# Proteogenomic reconstruction of organ-specific metabolic networks in an environmental sentinel species, the amphipod *Gammarus fossarum*

**DOI:** 10.1101/2024.05.04.592373

**Authors:** Natacha Koenig, Patrice Baa-Puyoulet, Amélie Lafont, Isis Lorenzo-Colina, Vincent Navratil, Maxime Leprêtre, Kevin Sugier, Nicolas Delorme, Laura Garnero, Hervé Queau, Jean-Charles Gaillard, Mélodie Kielbasa, Sophie Ayciriex, Federica Calevro, Arnaud Chaumot, Hubert Charles, Jean Armengaud, Olivier Geffard, Davide Degli Esposti

## Abstract

Metabolic pathways are targets of environmental contaminants underlying a large variability of toxic effects throughout biodiversity. However, the systematic reconstruction of metabolic pathways remains limited in environmental sentinel species due to the lack of available genomic data in many taxa of animal diversity. In order to improve the knowledge of the metabolism of sentinel species, in this study we used a multi-omics approach to reconstruct the most comprehensive map of metabolic pathways for a crustacean model in biomonitoring, the amphipod *Gammarus fossarum*.

We revisited the assembly of RNA-seq data by *de novo* approaches drastically reducing RNA contaminants and transcript redundancy. We also acquired extensive mass spectrometry shotgun proteomic data on several organs from *G. fossarum* males and females to identify organ-specific metabolic profiles.

The *G. fossarum* metabolic pathway reconstruction (available through the metabolic database GamfoCyc) was performed by adapting the genomic tool CycADS and we identified 377 pathways representing 7,630 annotated enzymes, 2,610 enzymatic reactions and the expression of 858 enzymes was experimentally validated by proteomics. Our analysis shows organ-specific metabolic profiles, such as an elevated abundance in enzymes involved in ATP biosynthesis and fatty acid beta-oxidation indicative of the high-energy requirement of the gills, or the key anabolic and detoxification role of the hepatopancreatic caeca, as exemplified by the specific expression of the retinoid biosynthetic pathways and glutathione synthesis.

In conclusion, the multi-omics data integration performed in this study provides new resources to investigate metabolic processes in crustacean amphipods and their role in mediating the effects of environmental contaminant exposures in sentinel species.

## Introduction

Metabolic pathways are potential targets of environmental contaminants, both in humans and in other species (Fritsche et al., 2023; Jordão et al., 2015; Massart et al., 2022). Recent studies showed that some chemicals, such as tributyltin, juvenoid hormones, or bisphenol A, can target and act as endocrine disruptors on lipid metabolism in the model species *Daphnia magna* (Fuertes et al., 2019; Jordão et al., 2016a, 2016b, 2015). These compounds have been shown to affect lipid (e.g. triacylglycerols and cholesterol) distribution, storage, or biosynthesis. Transcriptomic studies have also shown that changes following lipid accumulation include up-regulation of genes involved in fatty acid, glycerophospholipid and glycerolipid metabolism, membrane constituents, and chitin and cuticle biosynthesis pathways (Fuertes et al., 2019).

Although the use of a handful of model species plays a key role in discovering and acquiring knowledge on the molecular mechanisms underlying invertebrate biology and physiology, some obstacles, such as the phylogenetic distance with the species present in the diverse environments to be preserved, hampers their use in assessing the impact of environmental contaminants. Moreover, to improve environmental risk assessment, it is important to consider the biological diversity of a greater number of test species (Ruivo et al., 2022; Santos et al., 2018). In particular, metabolic networks (e.g. hormone synthesis pathways) show strong heterogeneity across phyla (Markov et al., 2009). Thus, strengthening the knowledge of the molecular physiology of sentinel species is an essential step to improve the extrapolation of toxicological effects of environmental contaminants across species. Unfortunately, there is a dramatic lack of genomic data, particularly in aquatic invertebrates used as sentinel species. To overcome this problem, the integration of mRNA sequencing with high-resolution mass spectrometry proteomics, an approach known as proteogenomics, has allowed the establishment of catalogs of thousands of proteins in a few of them, i.e. *Gammarus fossarum* or *Dreissena polymorpha* (Gouveia et al., 2018; Leprêtre et al., 2019; Trapp et al., 2014). Transcriptomic resources for gammarids typically include the pooling of RNAs coming from various individuals (Caputo et al., 2020; Trapp et al., 2014), but recent transcriptomes obtained from different gammarid species were obtained from genotyped single individuals, reducing the risk of chimeric transcript reconstruction (Cogne et al., 2019; Neuparth et al., 2020). In particular, our group contributed to those studies providing individual transcriptomes from individual male and female gammarids that were assembled separately to maximize mass spectrometry data extraction (Cogne et al., 2019). In this context, it would be of great interest to perform a *de novo* assembly using the RNA-seq data from both male and females organisms in order to improve gene coverage, to have a unique gene set for both sexes and to investigate potential contamination of the organism’s RNA from RNAs issued by its microbiome or by potential manipulation bias due to organisms’ size (a few centimeters). Similarly, metabolomics and lipidomics approaches in sentinel species such as the crustaceans *Gammarus. fossarum* or *Palaemon serratus* are showing species-specific lipid and metabolite compositions (Fu et al., 2021; Marie et al., 2023). Despite improvements in proteome and transcriptome functional annotations in the last decade, it has been noted that large discrepancies may occur in ontology assignment, especially in certain arthropod orders (McCartney et al., 2023), due to a lack of standardized and systematic bioinformatics methods.

Besides, most molecular studies in aquatic invertebrates used in ecotoxicology are carried out on whole-body sentinel organisms or pools of organisms (Besse et al., 2013; Kunz et al., 2010). This approach may help the identification of toxicity biomarkers (Leprêtre et al., 2022), but, as we have shown, the integration of organ-specific –omics profiles in emergent animal models is very relevant to improve our understanding of the molecular mechanisms (Degli Esposti et al., 2019; Koenig et al., 2021). For instance, the use of coexpression network analyses showed species-specific and orphan proteins involved in gonad maturation and embryonic development (Degli Esposti et al., 2019), or the mechanisms of toxicity of reproductive contaminants (Koenig et al., 2021) in *G. fossarum*. Organ gene expression analysis of newly identified metallothioneins transcripts showed a specific interaction between hepatopancreatic caeca and gills in the detoxification of heavy metals, such as Cadmium (Degli Esposti et al., 2024).

In this context, the present work aimed to provide a complete mapping of the metabolism of *G. fossarum* by adapting the Cyc Annotation Database System (CycADS), particularly suitable for metabolic gene annotation and network reconstruction using genomic data (Vellozo et al., 2011) to the use of transcriptomic data in the freshwater amphipod *G. fossarum.* Moreover, we aimed to integrate new and available proteomic datasets (Leprêtre et al., 2023) to identify organ-specific metabolic profiles and provide the basis to improve the knowledge of the effects of aquatic contaminants on metabolic pathways of this species and other closely related amphipods.

## Materials and methods

### Experimental design

To perform a first metabolic reconstruction based on the transcriptomic data available for *G. fossarum* (Cogne et al., 2019), we performed a new *de novo* assembly combining male and female RNA-seq raw data to reduce transcript redundancy and allow the implementation of CycADS. Subsequently, to integrate mRNA and protein expression levels, proteomic data from male and female organs of *G. fossarum* were obtained by label-free high-resolution mass spectrometry. Transcriptomic and proteomic data were integrated by cross-referencing the respective EC numbers to validate the expression of the enzymes identified and the proteomic dataset was further analyzed to investigate organ-specific metabolic pathway expression (Figure 1).

**Figure 1.**
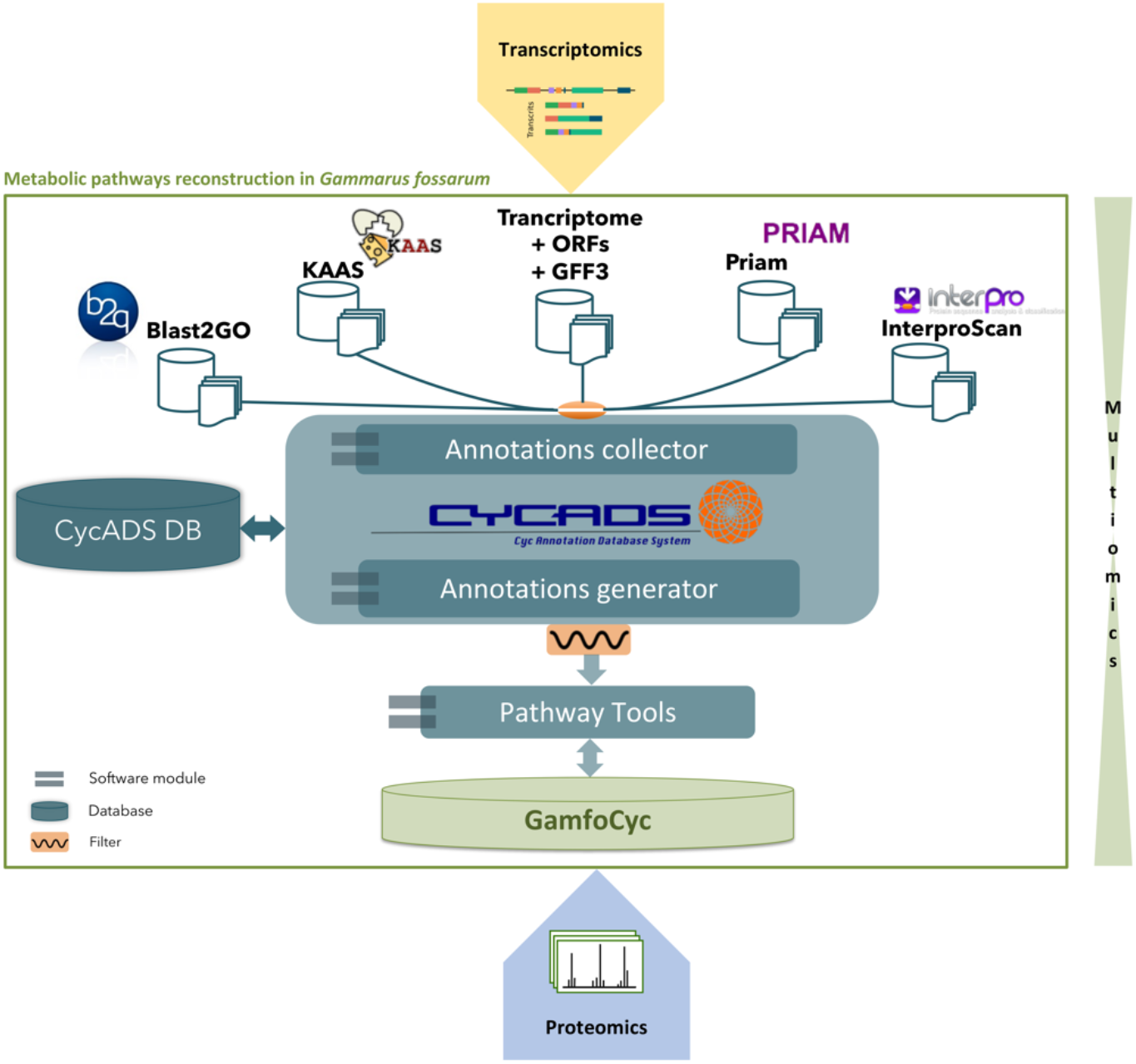
Outline of the methodological approach for the characterization of global metabolism in *Gammarus fossarum* using multi-omics integration involving transcriptomics, proteomics, and the CycADS annotation management system (adapted from (Vellozo et al., 2011). All software were used with their default settings, except for Blast2GO (e-value 10-5 against Swiss-Prot reference database), Interproscan (all sub-methods), Kegg (2 references « for genes » and « for eukaryote »). All annotation were collected by CycADS then extracted without any filter.

### Transcriptomic resources

RNA-seq data were retrieved from Cogne et al. (2019). Male and female gammarid data belonged to genotyped individuals of the species *G. fossarum*, subtype B. Illumina TruSeq stranded mRNA Sample Prep kit was used to generate cDNA libraries from poly-A enriched RNAs for each of the samples, which were then sequenced on two lanes of Hiseq3000 (Illumina) using a paired-end read length of 2 × 150bp with the HiSeq Reagent Kits (Illumina, San Diego, California) (Cogne et al. 2019). The raw reads are available in the “NCBI Sequence Reads Archive” under accession number SRR8089722 (“RNAseq GFBM,” 2018) (BioProject PRJNA497972 and BioSample SAMN10259937) for the male and under accession number SRR8089729 (“RNAseq GFBF,” 2018) (BioProject PRJNA497972 and BioSample SAMN10259936) for the female.

### De novo transcriptome assembly

#### Pre-processing of RNA-Seq data

A FastQC v0.11.9 analysis of the RNA-seq raw read dataset was performed (Figure S1, Pre-processing step 1), for all forward and reverse reads (male and female samples). Forward reads from male and female samples were merged in a single file, as well as reverse reads, to pool the RNA-seq data in a single paired-end dataset. The mean quality scores, defined as the mean quality value across each base position in the read, were satisfying for every sample (Phred > 28). All sequence lengths were 150 bp (base pair) long.

Trimmomatic v0.36 (Bolger et al., 2014) was used to trim residual adapter sequences and/or low-quality bases (Figure S1, Pre-processing, step 2), with palindrome mode and a PHRED score threshold set to 5 to avoid excessive trimming, according to MacManes’ benchmarking (2014). Only paired reads were kept.

To reduce RNA contamination possibly coming from organisms’ microbiome and sample manipulation, we identified nucleic sequences using the taxonomic sequence classifier Kraken2 v2.1.2 (Wood et al., 2019; Wood and Salzberg, 2014) (Figure S1, Pre-processing, step 3). Taxon information was obtained from the NCBI taxonomy database (Federhen, 2012). Taxonomic classification was run using a complete set of reference libraries, comprising RefSeq complete sets of proteins from Archaea, Bacteria, Fungi, Plant, and Protozoa taxa as well as plasmid and viruses and GRCh38 human proteins (O’Leary et al., 2016).

To further exclude from the raw reads any rRNA sequence residues, SortMeRNA v4.3.4 (Kopylova et al., 2012) was used for filtering rRNA sequences using SILVA database (Quast et al., 2013) with representative small and large subunit rRNA sequences from Archaea, Bacteria, and Eukarya (Figure S1, Pre-processing, step 4).

Last quality control was performed to ensure that data met assembly requirements and to retrieve pre-processing statistics with the MultiQC v1.12 tool (Ewels et al., 2016) (Figure S1, Pre-processing, step 5).

#### Assembly pipeline

*De novo* assembly of *G. fossarum* transcriptome was performed using the Trinity v2.14.0 pipeline (Grabherr et al., 2011; Haas et al., 2013) (Figure S1, Normalization and Assembly, step 1). We combined the standard procedure using the three independent modules (Inchworm, Chrysalis, and Butterfly) in addition to the *in silico* normalization procedure to reduce assembly time and errors.

A general assembly metrics report, including total trinity transcripts, median contig length, average contig, total assembled bases, etc. was obtained with a Trinity embedded script (https://github.com/trinityrnaseq/trinityrnaseq/blob/330b6fe9c65af0e203c5620708cce0fd6f57ceab/util/TrinityStats.pl). TransDecoder v5.5.0 (Haas, 2018) was used to identify candidate coding regions within transcript sequences (Figure S1, Normalization and Assembly, step 2). We estimated the completeness and redundancy of processed data with the Benchmarking Universal Single-Copy Orthologs (BUSCO) v5.3 (Simão et al., 2015), based on universal single-copy orthologs in the OrthoDB database (Figure S1, Normalization and Assembly, step 3) (Kriventseva et al., 2019). The *arthropoda_odb10* database, made up of 1,013 arthropod single-copy orthologs was used, as it is the closest taxonomically to gammarids. BUSCO was run in transcriptome mode with the default options.

To reduce the redundancy of alternative transcripts, we first chose to filter out possible transcript artifacts and transcripts with low expression levels as recommended by (Haas et al., 2013). Then, we evaluated the percentage of expression level (abundance) for a given transcript compared to all transcripts within an isoform cluster (Haas et al., 2013), the so-called isoform percentage (IsoPct), to retain the highest abundant isoform by applying the “*Highest Iso Only*” option (Figure S1, Normalization and Assembly, step 4). To compute transcript isoform abundances as normalized expression values (Fragments Per Kilobase of transcript per Million FPKM), the RSEM (Li and Dewey, 2011) and Bowtie2 v2.4.5 (Langmead and Salzberg, 2012) tools were used.

To assess the quality of the final assembly, we performed a new BUSCO analysis (Figure S1, Normalization and Assembly, step 5).

#### Functional annotation of reference transcriptome by CycADS and metabolic network reconstruction

CycADS v1.36 is a system that allows the standardization of genome annotation, executed by command line and customizable with a configuration file (Vellozo et al., 2011). In this study, in order to adapt a transcriptomic assembly for CycADS, we created a structural annotation file (GFF3) with TransDecoder v5.5.0 (Haas, 2018). Then, CycADS was used according to the methodology described by Vellozo et al. (2011) (Figure 1).

In particular, we performed a functional annotation on the assembled transcriptome with pipelines on local machine, namely Blast2GO v2.5 (Conesa et al., 2005; Conesa and Götz, 2008), UniProtKB/Swiss-Prot protein database (The UniProt Consortium, 2021), Priam v2_JAN_18 (Claudel-Renard et al., 2003), InterproScan v5.31-70.0 (Jones et al., 2014), and the KAAS-KEGG v2.1 online pipeline (Moriya et al., 2007). Functional information was collected (KEGG Orthology (KO), Enzyme Commission (EC) number, and Gene Ontology (GO)) from the annotation methods outputs using the CycADS annotation collector module (Figure 1). All annotations were then extracted and written in an enriched “Pathological file” (PF) which was used in the “Pathway Tools” compartment (Karp et al., 2010) to perform the metabolic reconstruction and generate the corresponding BioCyc Pathway/Genome Database (PGDB), named GamfoCyc (Figure 1). Software settings are given in Figure 1.

For the following analyses, to focus on the metabolic network, we chose to work only on the subset of enzymes involved in the metabolic pathways (see results and discussion part).

The MetExplore v2.30.8 platform (Cottret et al., 2018) has been developed to explore metabolic pathways, manipulate the metabolic network graph, and map omics data. Here, we used the online platform to analyze the completeness of each identified metabolic pathway. The network was therefore exported as SBML and BioPAX files containing gene products, genetic enzyme complexes, reactions, pathways, and metabolites and was then imported into MetExplore.

The GamfoCyc database was added to the Arthropodacyc metabolic database collection (available at http://arthropodacyc.cycadsys.org/) and the MetExplore database (available at https://metexplore.toulouse.inrae.fr/GAMFO_GFB).

#### Proteomic resources

Proteomics data were acquired from three males and three females of *G. fossarum* species. Six organs or anatomical regions (cephalon gills, caeca, intestine, male gonads, female gonads, and the rest of the body) were sampled and collected for each organism and frozen in liquid nitrogen before protein extraction. Gills and caeca proteomes were previously described in Leprêtre et al., 2023.

#### Protein extraction

Each organ was directly dissolved in 40 μL of LDS sample buffer (Invitrogen), except for the cephalon and the rest of the body, which were previously ground in LDS by adding a 4-mm steel ball and then using a tissue homogenizer. Samples were subjected to 1 min of sonication (transonic 780H sonicator) and were heated for 5 min at 95 °C. Organ shreds were completely dissolved in LDS sample buffer, and then 35 μL of each sample was subjected to SDS-PAGE on a 10-well 4-12% gradient NuPAGE (Invitrogen) for 10 min at 150 V with MES buffer. Gels were stained with Coomassie Blue Safe dye (Invitrogen) and destained overnight with water. The total protein in each well was extracted as a single polyacrylamide strip and processed for further decoloring and iodoacetamide treatment. Proteins were proteolyzed with sequencing grade trypsin (Roche) using 0.01% Protease-MAX surfactant (Promega).

#### Mass spectrometry

The resulting peptide mixtures were analyzed in data-dependent acquisition mode with an Orbitrap Exploris high-resolution mass spectrometer (MS) (ThermoFisher) coupled to an UltiMate 3000 LC system (Dionex-LC Packings), operated as described previously (Ramos-Nascimento et al., 2023).

#### Protein identification and quantification by spectral counting

Peak lists were generated with Mascot Daemon software (version 2.3.2; Matrix Science) using the data import filter extract_msn.exe (Thermo). Data import filter options were set to 400 (minimum mass), 5,000 (maximum mass), 0 (clustering tolerance), 0 (intermediate scans), and 1,000 (threshold), as described previously (Christie-Oleza et al., 2012). MS/MS spectra were assigned to peptide sequences with the Mascot Daemon 2.3.2 search engine (Matrix Science) against the custom database derived from the assembled transcriptome. Protein detection was validated with at least 1 specific peptide and 2 peptides in total.

Two tables were obtained for each individual from mass spectrometry. The first table (Table MS1 available at https://doi.org/10.57745/HMQVCS) corresponds to the peptides (32,925 unique male peptide sequences, 39,018 unique female peptide sequences) detected by mass spectrometry and their associated characteristics (associated protein identifier, sequence, length, mass/charge (*m/z*) ratio, modifications, retention time, Mascot score, etc.). The second table (Table MS2, available at https://doi.org/10.57745/HMQVCS) contains the protein abundance measurements for each sample expressed as “spectral count” (SC). Indeed, spectral counts are an estimate of the abundance of a protein in the samples through the number of MS/MS spectra that are associated (Liu et al., 2004). The more abundant a protein is, the more likely its peptides will be frequently fragmented in the mass spectrometer. For the male samples, we have a matrix of 5,073 identified proteins (Table MS2 GFBM). For the female samples, we have a matrix of 5,678 identified proteins (Table MS2 GFBF).

#### Proteomic data integration

We retrieved EC numbers from transcriptomic annotation (GamfoCyc) (n=4,033) and EC numbers of proteins identified in shotgun proteomics (all organs combined) (n=4,150; available at https://doi.org/10.57745/HMQVCS). These two lists of ECs were mapped onto the metabolic maps of several pathways in the KEGG database (Kanehisa and Goto, 2000) through to the KEGG Mapper collection of tools (Kanehisa et al., 2022; Kanehisa and Sato, 2020). This approach enables us to observe the ability of shotgun proteomics to confirm the presence of putative proteins found in transcriptomics data.

To retain proteins whose abundance may be low and specific to an organ and exclude those whose detection is incidental, only proteins identified by at least one spectral count and present in at least 3 samples were retained (Gregori et al., 2013). From the filtered table, differential analysis of protein abundance in organs was performed using the R package EdgeR (Robinson et al., 2010) version 3.32.1.

Although this function was originally designed for RNA-Seq count data, it is also applicable to spectral count data in proteomics (Gregori et al., 2013). The selection of differentially expressed (DE) proteins was based on a False Discovery Rate (FDR) threshold of 0.05 and an absolute change in expression of 2 (|Fold Change| > 2) (Table S2).

#### Pathway analysis

We collected the list of DE proteins found in abundance in gills (n=653), caeca (n=635), male gonads (n=343), and female gonads (n=187), and used them as an input list for the pathway enrichment analysis. We performed an overrepresentation analysis (ORA) (Khatri et al., 2012) from the WebGestalt server (Liao et al., 2019), using the KEGG pathways databases (Kanehisa et al., 2022). The tool was used with the following options: FDR (Benjamini-Hocheberg adjusting method), at least three genes from the input list in the enriched category, and the whole potential coding transcriptome (n=63,639 ORFs) as the reference background. We obtained an enrichment ratio (ER), which is the number of observed proteins divided by the number of expected proteins from each KEGG category in the n-protein list (Liao et al., 2019). We thus obtained a percentage of proteins overrepresented in the pathway of interest. We chose to use the “weighted set cover” method to reduce the redundancy of the gene sets in the enrichment result to identify the most representative significant gene sets for visualization (Liao et al., 2019).

## Results and discussion

### A reference transcriptome for the sentinel species Gammarus fossarum (subtype B)

In this work, we pooled female and male individual RNA-seq datasets from *G. fossarum* (subtype B) individuals to obtain a new reference transcriptome for this species.

The number of merged male and female reads is 181,487,648 (around 90.7 M forward and 90.4 M reverse) (Table 1). We identified 23,361,906 reads (12.9%) (Table 1) as belonging to different taxa using Kraken 2 (Wood et al., 2019; Wood and Salzberg, 2014) and we identified bacterial RNAs as the main contaminant source (5.34%), followed by green plants RNAs (2.05%) and fungi (1.1%). *Homo sapiens* RNAs represented the single species main contaminant source (2.05%) (Figure S2A). In parallel, the search for rRNAs with SortMeRNA (Kopylova et al., 2012) highlighted 674,405 reads (0.37%) coming from ribosomal databases (Table 1), of which 63,230 matched a hit in the SILVA archaeal database (Quast et al., 2013), 110,652 in the bacterial database, and 474,224 in the eukaryotic database (Figure S2B). The total number of read after the contamination removal with Kraken2 and SortMeRNA amounted at 157,400,267 (86.73%) and were used for the *de novo* assembly of *G. fossarum* transcriptome and its metabolism reconstruction.

**Table 1.**
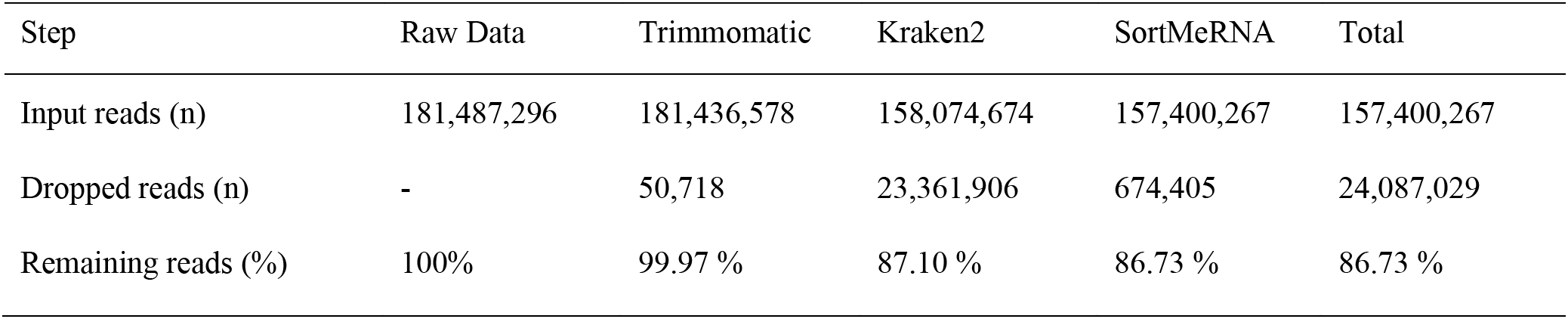
Summary of the pre-processing results.

| Step | Raw Data | Trimmomatic | Kraken2 | SortMeRNA | Total |
| --- | --- | --- | --- | --- | --- |
| Input reads (n) | 181,487,296 | 181,436,578 | 158,074,674 | 157,400,267 | 157,400,267 |
| Dropped reads (n) | - | 50,718 | 23,361,906 | 674,405 | 24,087,029 |
| Remaining reads (%) | 100% | 99.97 % | 87.10 % | 86.73 % | 86.73 % |

The results of our pre-processing step show that RNA contamination from food sources, organisms’ microbiomes, and even experimental manipulation must be considered to limit artifacts of transcriptome assembly of novel species. To our knowledge, this has not been done for most environmental species (Caputo et al., 2020; Cogne et al., 2019; Llorente et al., 2020). In parallel, these results show the interest of considering RNA contaminants to reconstruct the holotranscriptome (host transcriptome and microbiota transcriptome) of *G. fossarum*, similarly to recent hologenome approaches performed in *Daphnia magna* or the metaproteomic studies recently tested in gammarids (Chaturvedi et al., 2023, Gouveia et al 2020).

Since the initial assembly strategy of the two original transcriptomes (Cogne et al., 2019) may involve the potential presence of redundant transcripts and isoforms, we took into account the expression level of the transcripts and to keep only the most highly expressed transcript (i.e., the highest isoform) in the refined final assembly (Haas et al., 2013). After excluding lowly expressed isoforms with the RSEM tool, the refined reference transcriptome contained 71.10% (302,024) of the original transcripts (Table 2). Compared with the primary assembly the number of transcripts under 200 bases decreased to close to zero in the filtered assembly (“n_under_200” from 49 to 2, Table 2). The percentage of contigs with an ORF is higher in the new transcriptome assembly (“mean_orf_percent” from about 48% to about 54%, Table 2).

**Table 2.**
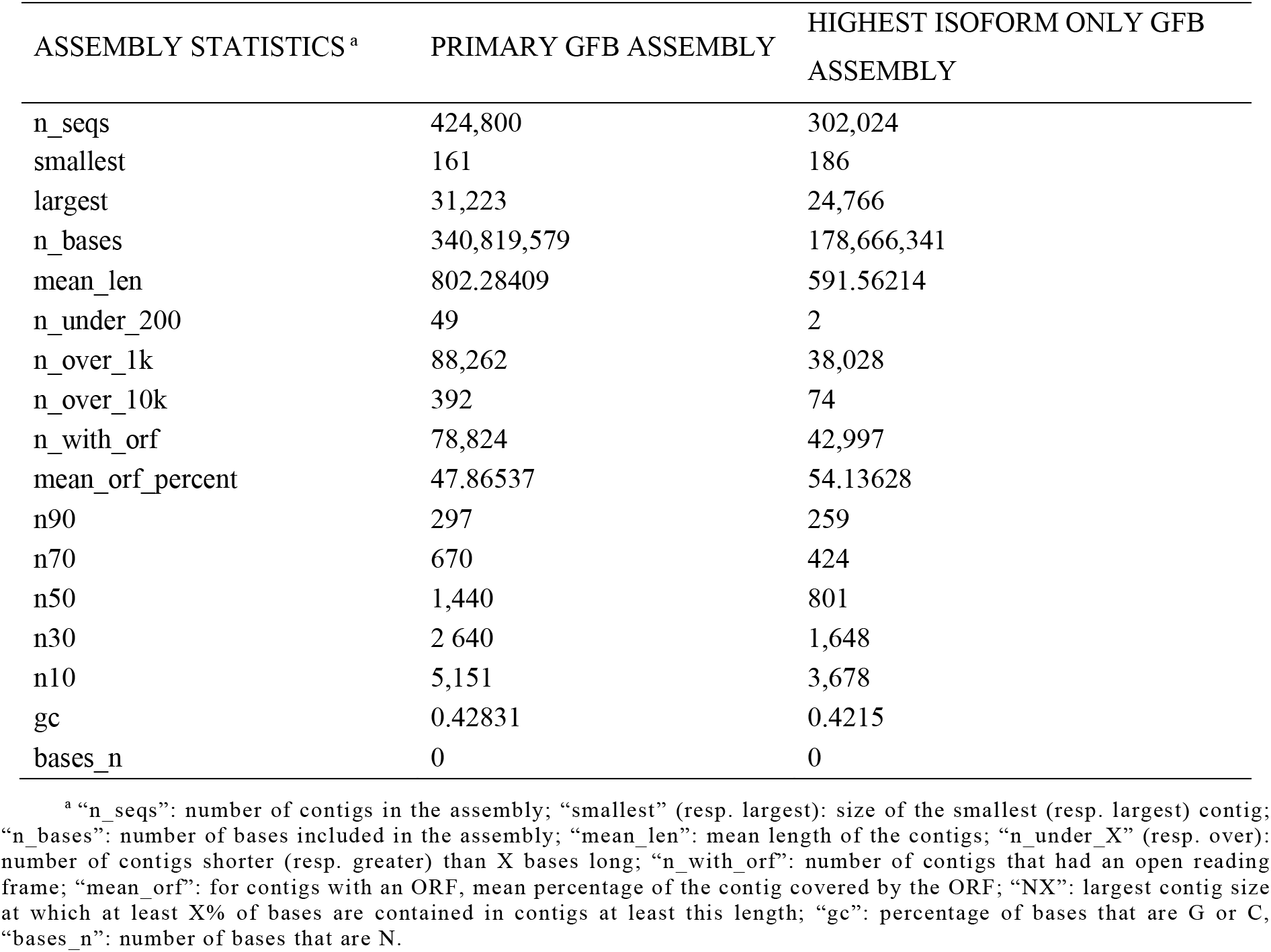
Metrics for the *Gammarus fossarum* transcriptome assemblies.

| ASSEMBLY STATISTICS<sup>a</sup> | PRIMARY GFB ASSEMBLY | HIGHEST ISOFORM ONLY GFB ASSEMBLY |
| --- | --- | --- |
| n_seqs | 424,800 | 302,024 |
| smallest | 161 | 186 |
| largest | 31,223 | 24,766 |
| n_bases | 340,819,579 | 178,666,341 |
| mean_len | 802.28409 | 591.56214 |
| n_under_200 | 49 | 2 |
| n_over_1k | 88,262 | 38,028 |
| n_over_10k | 392 | 74 |
| n_with_orf | 78,824 | 42,997 |
| mean_orf_percent | 47.86537 | 54.13628 |
| n90 | 297 | 259 |
| n70 | 670 | 424 |
| n50 | 1,440 | 801 |
| n30 | 2,640 | 1,648 |
| n10 | 5,151 | 3,678 |
| gc | 0.42831 | 0.4215 |
| bases_n | 0 | 0 |

<sup>a</sup> “n\_seqs”: number of contigs in the assembly; “smallest” (resp. largest): size of the smallest (resp. largest) contig; “n\_bases”: number of bases included in the assembly; “mean\_len”: mean length of the contigs; “n\_under\_X” (resp. over): number of contigs shorter (resp. greater) than X bases long; “n\_with\_orf”: number of contigs that had an open reading frame; “mean\_orf”: for contigs with an ORF, mean percentage of the contig covered by the ORF; “NX”: largest contig size at which at least X% of bases are contained in contigs at least this length; “gc”: percentage of bases that are G or C, “bases\_n”: number of bases that are N.

To evaluate the impact of the pre-processing strategy as well as the decrease in redundancy of isoforms in our filtered assembly, we performed a BUSCO analysis. The BUSCO results show that more than 92% of the 1,013 *Arthropoda* orthologous genes (sum of complete and fragmented) are found in the primary transcriptome, and 90% in the refined transcriptome (Figure 2). This indicates that our refined *G. fossarum* transcriptome assembly did not lose much of the expected biological information from its phylogenetic membership. About 38% (381 out of 1,013) of the complete genes were found in a single copy in the primary transcriptome, against about 63% (638 out of 1,013) in the refined transcriptome (Figure 2). Thus, our pipeline applied to the RNA-seq data led to an increased presence of single-copy genes (+ 25%) and a decreased number of duplicated orthologs (-28%), with only a slight increase (of about 2%) in the number of missing genes. These results suggest that using a biology-driven approach that takes into consideration RNA origins and level of expression may increase the species-specificity of *de novo* assemblies. The remaining duplicated genes in the refined transcriptome could come either from different haplotypes or isoforms still present in the assembly or from true duplications in the amphipod family, a hypothesis that should be tested using new genomic resources. Additionally, it must be noted that the BUSCO arthropod database derives from OrthoDB, which contains only *Daphnia pulex* as a crustacean species (Kriventseva et al., 2019), indicating an under-representation of crustaceans in the ortholog database. The completeness of the transcriptomes is positively correlated to the proportion of complete BUSCO genes, but this evaluation may be biased by the number of species close to the species of interest (Amil-Ruiz et al., 2021; Seppey et al., 2019). In summary, this new assembly provides a common annotation of *G. fossarum* male and female organisms with improved transcript coverage and reduced redundancy and external RNA contamination.

**Figure 2.**
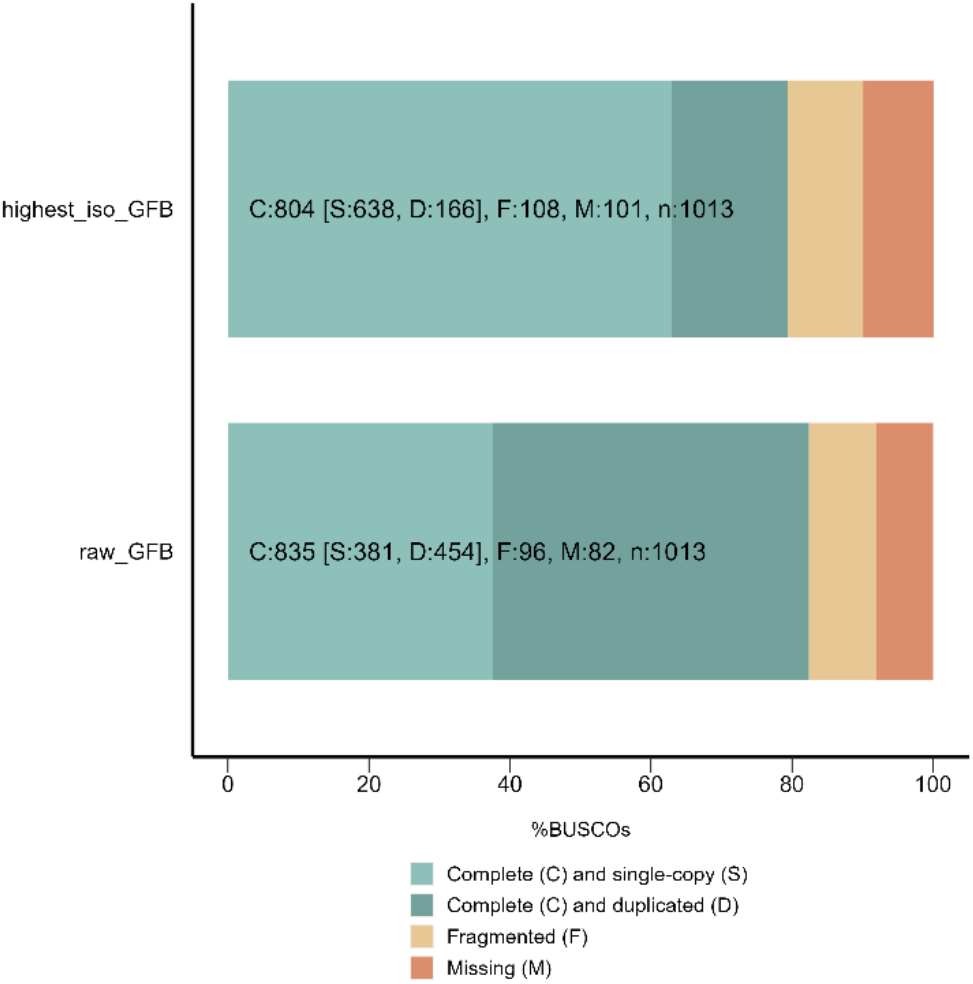
Completeness analysis of the pre-filtered (raw_GFB) and filtered reference (highest_iso_GFB) transcriptomes of *G. fossarum* via BUSCO of arthropods.

### Development of the metabolism database (GamfoCyc) dedicated to Gammarus fossarum

The refined transcriptome was then annotated by the CycADS pipeline (Vellozo et al., 2011) to create a database of the metabolic pathways of *G. fossarum*. This database, named GamfoCyc, was obtained from the analysis of 63,639 polypeptides/ORFs sequences (available at https://doi.org/10.57745/HMQVCS) from 42,997 contigs (Table 2). Four methods were used to conduct the functional annotation of the *G. fossarum* transcriptome, collecting 7,630 enzymes classified as complete, i.e. EC numbers with all the 4 levels of enzyme classification defined representing 1,764 distinct EC numbers (Figure 3A). Of these 7,630 enzymes, 4,033 are involved in the metabolic pathways identified by BioCyc and thus constitute the *G. fossarum* metabolic network. All functional analyses were therefore performed on this subset of 4,033 enzymes. KAAS and BLAST2GO were the major sources of annotation with 1,332 and 1,166 annotations respectively, compared to Interpro (447 annotations) and Priam (412 annotations) (Figure 3A). There were 194 EC numbers common to all methods (Figure 3A). We can also note that 840 EC numbers are not cross-referenced, which is due on the one hand to the different annotation strategies that do not target the same features (i.e., local alignment vs sequence profile-based searches or functional domain searches) and on the other hand to the incomplete EC numbers (i.e., not all classification levels are defined) which are not taken into account by PathwayTools (Figure 3A).

**Figure 3.**
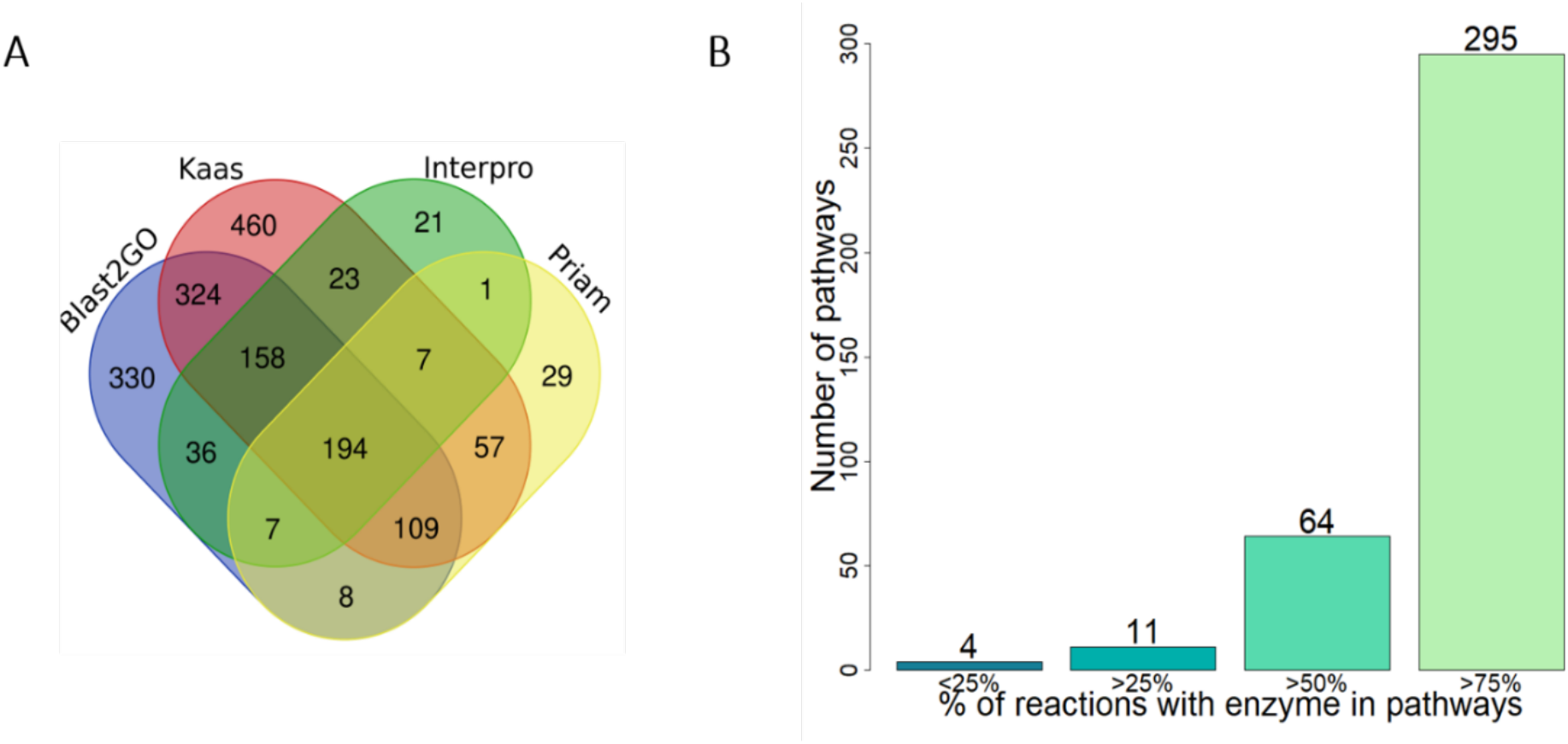
Functional annotation of the reference transcriptome ( A) Venn diagram of EC annotation in *Gammarus fossarum* according to the different methods (Blast2GO, KAAS-KEGG, InterproScan, Priam) of the CycADS pipeline (B) Completeness analysis of metabolic pathways annotation via MetExplore.

The collection of functional annotations allowed the reconstruction of metabolic pathways, and the mapping of 7,630 enzymes onto these pathways (Table S1). Our filtering method does not identify annotation artifacts (e.g., transcripts may be incomplete). In fact, from a biological point of view, if two isoforms are annotated as conducting the same enzymatic reaction, the Pathway Tools tool keeps both annotations. In this case, the presence of true isozymes is also possible.

In order to assess the network quality by measuring the completeness of the metabolic pathways in GamfoCyc, the MetExplore platform (Cottret et al., 2018) was used to assess the percentage of reactions annotated with enzymes in pathways. A comparison with the metabolic network of two model organisms, *Drosophila melanogaster* and *Daphnia pulex,* was also performed. The results showed that the GamfoCyc database contains 374 annotated metabolic pathways composed of 2,610 reactions (Table S1). Among the annotated metabolic pathways, 295 pathways (79 %) contained more than 75% of reactions annotated with enzymes. Only 4 pathways (1%) contained less than 25% of annotated reactions (Figure 3B).

These latter are involved in eumelanin biosynthesis, iron-sulfur cluster biosynthesis, protein nitrosylation/denitrosylation, and glycosphingolipids biosynthesis. Interestingly, glycosphingolipids are rarely identified in the lipidome studies on gammarids (Arambourou et al., 2018; Fu et al., 2021) and our metabolic reconstruction sheds light on these previous findings suggesting that this pathway is either absent or not detectable at gene expression level in *G. fossarum*. In the case of eumelanin biosynthesis, scarce knowledge of this metabolic pathway in *Arthropoda* may explain this result, since arthropod cuticular melanin has been shown to be different from mammalian epidermal melanins (Barek et al., 2018).

We therefore compared our results with data from these species to illustrate the relevance of our strategy for reconstructing the metabolic pathways of a non-model organism from the transcriptome. As a comparison, in the *Drosophila melanogaster* database (genome version release 6 plus ISO1MT, (Hoskins et al., 2015), 4,965 enzymes are annotated and in the *Daphnia pulex* database (genome version jgi_v11 geneset, (Colbourne et al., 2011) 3,672 enzymes are annotated (Baa-Puyoulet et al., 2016; Consuegra et al., 2020; McQuilton et al., 2012, Nordberg et al., 2014). As both species have annotated genomes, the large number of enzymes found in *G. fossarum* (n=7,630) from the transcriptomes could be explained by the presence of multiple annotated transcripts for the same enzyme, despite the reduced redundancy of our assembly. The reconstructed global metabolism of *D. melanogaster* contains 2,201 reactions for 353 pathways (Figure S3). Regarding the completeness of the annotation of the globality of the pathways, 277 pathways (78.4%) contain more than 75% of reactions with annotated enzymes, and 5 pathways (1.4%) contain less than 25% of reactions with annotated enzymes (Figure S3). Thus, the results obtained for *G. fossarum* transcriptome are qualitatively similar to those of a model organism for which the reconstruction of the global metabolic pathways was performed using a sequenced and annotated genome. This work shows that the exploitation of transcriptomic data is therefore feasible and relevant in a non-model species by adapting the CycADS tool to this type of data and opens new perspectives for other non-model species for which data are already available.

### Integration of proteomic data for metabolic annotation

After the reconstruction of *G. fossarum* metabolic pathways from the transcriptomic data, we validated the presence of the enzymes and studied the organ-specific metabolic profiles by integrating the shotgun mass spectrometry proteomic data from six distinct organs or anatomical regions (cephalon, gills, caeca, intestine, gonads, and the rest of the body) sampled individually from three male and three female *G. fossarum B* individuals. In female organs, we obtained 35,515 peptides of different sequences and 3,643 proteins and a cumulative 466,942 spectral counts for all 18 samples analyzed (Figure S4A). In male organs, we obtained 29,174 peptides of different sequences and 3,133 proteins (validated with at least one specific peptide and 2 peptides in total) and cumulatively 353,123 spectral counts for all 18 samples analyzed (Figure S4A). By cross-referencing male and female tables of spectral counts, the overall proteomic profile of *G. fossarum* was composed of 4,150 proteins in total. This refined version of the *G. fossarum* reference transcriptome allowed for increasing the proteomic catalog of this species by more than two-fold compared to the first reported proteogenomic study (n=1,873) (Trapp et al., 2014).

In total, 858 enzymes annotated in the metabolic pathways were experimentally validated by mass spectrometry measurements. This represents 21.3% of the enzymes mapped on the transcriptome (n=4,033) and 20.7% of the proteins identified by shotgun proteomics (n=4,150) (Figure S4).

To find and visualize where enzymes are involved in metabolic reactions, we chose to map the enzymes annotated by the transcriptome and validated in proteomics onto the metabolic pathways. As an example, the mapping of the enzymes validated by proteomics on the beta-oxidation pathway (more than 75% of the reactions annotated) is shown in Figure 4. In contrast, the glycosphingolipid biosynthesis pathway, (less than 25% reactions annotated) showed only two enzymes identified by shotgun proteomics (Figure S5).

**Figure 4:**
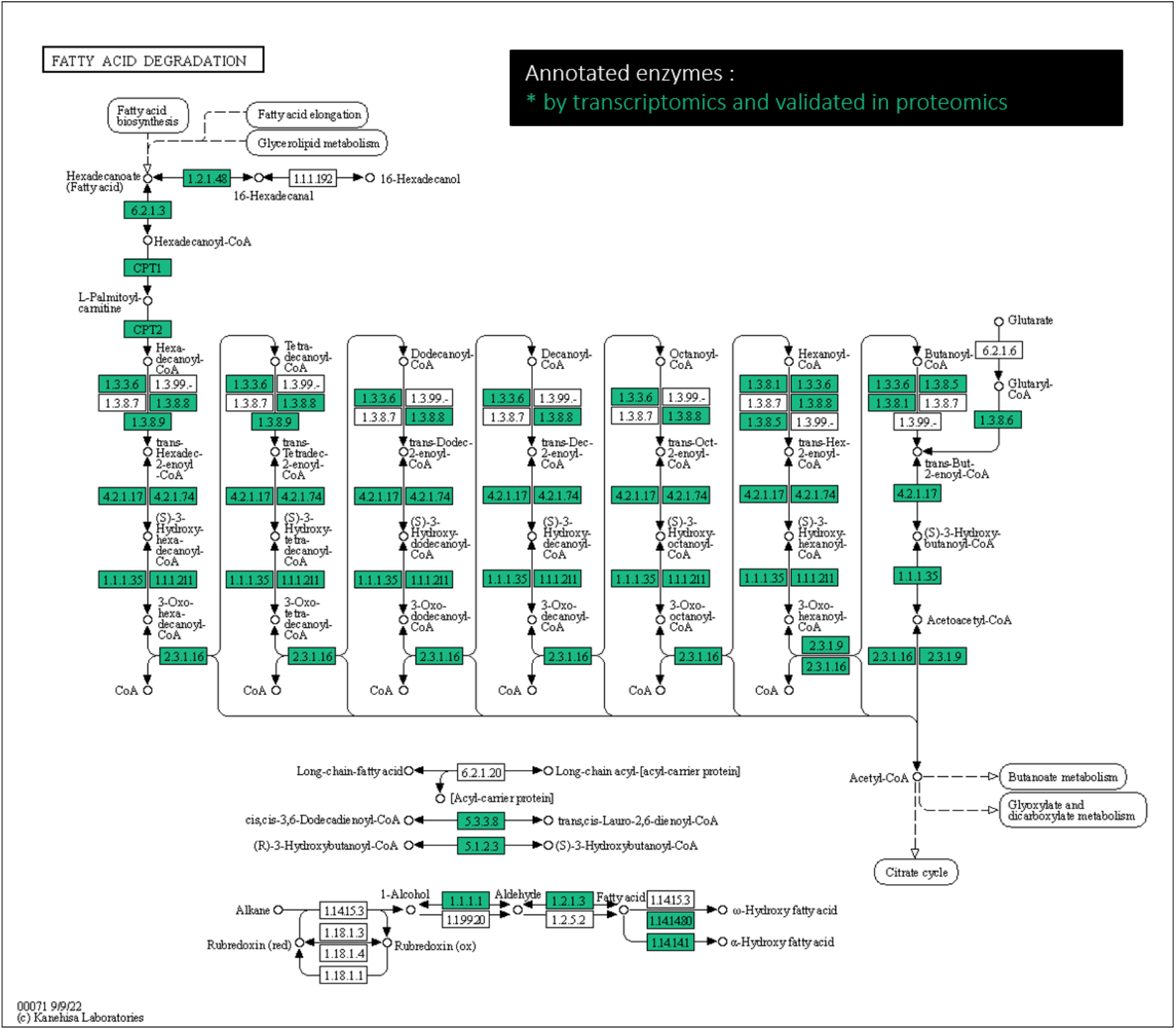
Example of mapped enzymes of *Gammarus fossarum* involved in the beta-oxidation pathway. Pathway module (functional unit of gene sets in metabolic pathway) is highlighted in green when the correspondent EC was annotated by transcriptomics and validated in proteomics.

The integration of proteomic data that we proposed in this work provides the first multi-omics and experimental validation of metabolic pathways in an environmental sentinel species.

### Organ-specific metabolism through proteomic profiling

We retained 2,190 proteins across the proteomic dataset to investigate the organs-specific basal metabolism in gammarids, after filtering out lowly abundant proteins and spectral count normalization.

A multivariate analysis of the data shows organ-specific protein profiles is shown in Figure 5. Proteome profiles showed an expected sexual dimorphism in the gonads, and a general weak variability in biological replicates (Figure 5). In fact, inter-sex comparisons in each single organ showed few significantly differentially expressed proteins. For example, male and female gills differed by around 1% (n=21) of their proteome (Figure S6). Similar results were found for the caeca, approximately, with 1.6% (n=35) of proteins found differentially expressed between male and females (Figure S7). These results suggest a low inter-sex variability between *G. fossarum* organs at the proteome level, except for the gonads (Figure 5). Intestines showed higher individual variability compared to other organs, probably due to a lower protein content (mirrored by a lower spectral count compared to other organs) in intestine protein extracts, and therefore a more difficult reproducibility for these organs.

**Figure 5.**
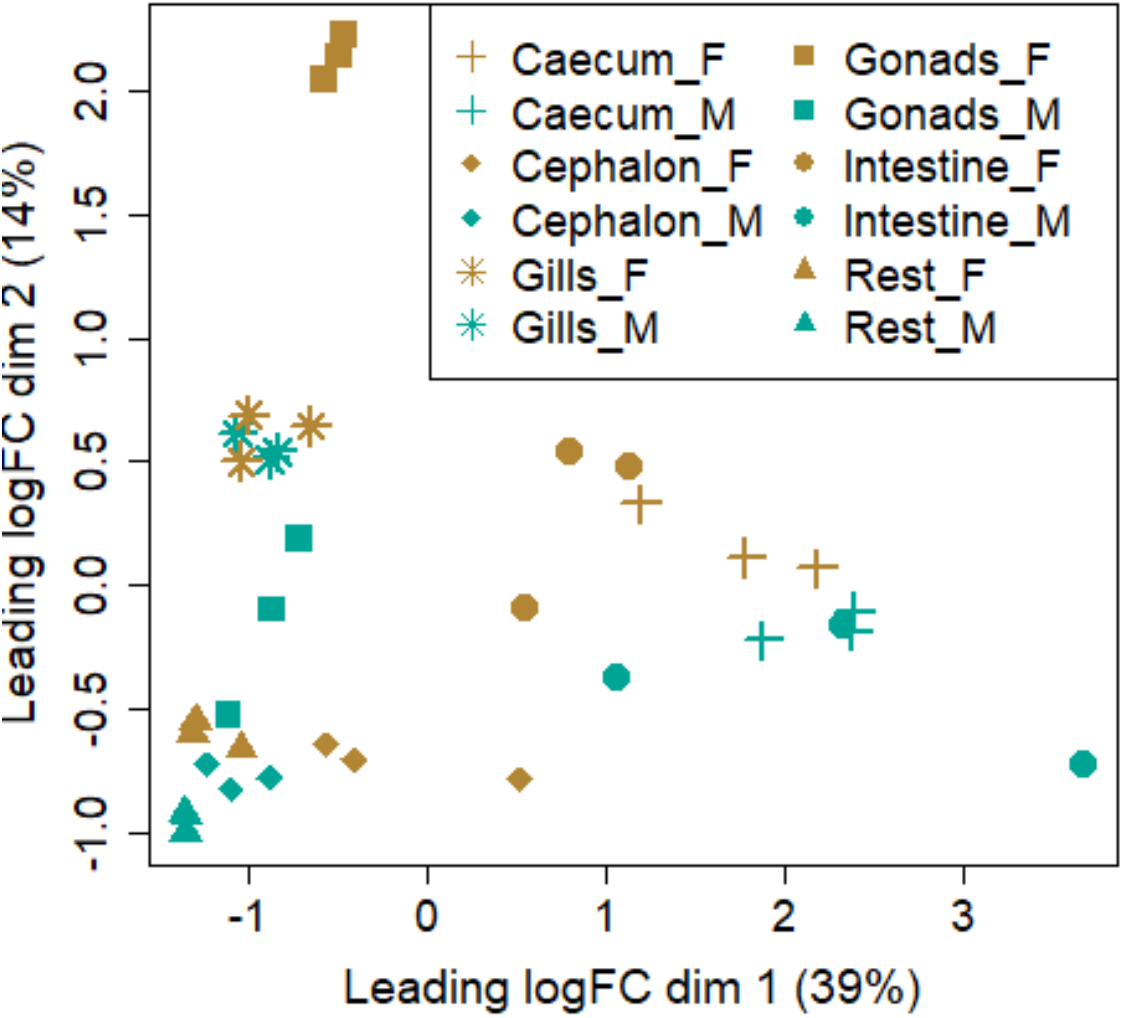
: Multi-dimensional scaling (MDS) plot of expression profiles of male (*_M) and female (*_F) samples in *Gammarus fossarum* proteome.

In order to investigate organ-specific metabolic profiles, we first compared the proteomes and the metabolic pathways in the gills and the caeca compared to the other organs (Figure 6, Table S2). In the gills we found 653 proteins more abundant than the other organs (FDR<0.05, FC>2), of which 237 are annotated in GamfoCyc (Figure 6A, Table S2). Proteins involved in energy metabolism were predominant in the gills, in particular the pathways of the TCA cycle with an ER of 30.17% (FDR<2.2e-16) (Figure 6B, Table S5). Proteins involved in oxidative phosphorylation and the degradation of fatty acids were also enriched, with an enrichment of around 20% (FDR <2.2e-16) and 16% (FDR=1.58e-10), respectively (Table S5). Similar results have been shown previously in gammarids (Leprêtre et al., 2023) and may be explained by the extremely energy demanding processes the gills are responsible for, such as oxygen uptake, acid–base balance, and osmotic and ionic regulation (Henry and Wheatly, 1992; Péqueux, 1995). As an example, osmoregulation is suggested to represent 11% of the total energy budget in *Gammarus pulex* (Felten et al., 2008). On the other hand, oxidative phosphorylation is the most effective energy-release process in animals (Dimroth et al., 2000; Jin et al., 2019). This pathway also appears to be modulated by abiotic conditions such as salinity in aquatic organisms (Bal et al., 2021), and pollutants in *Gammarus* spp. (Kunz et al., 2010) or plays a role in the adaptation of *Gammarus lacustris* to environmental conditions in the Tibetan region (Jin et al., 2019). Altogether, these results point out a key role of cellular respiration based on ATP biosynthesis and fatty acid consumption as a major energy resource in the gills of *G. fossarum*.

**Figure 6.**
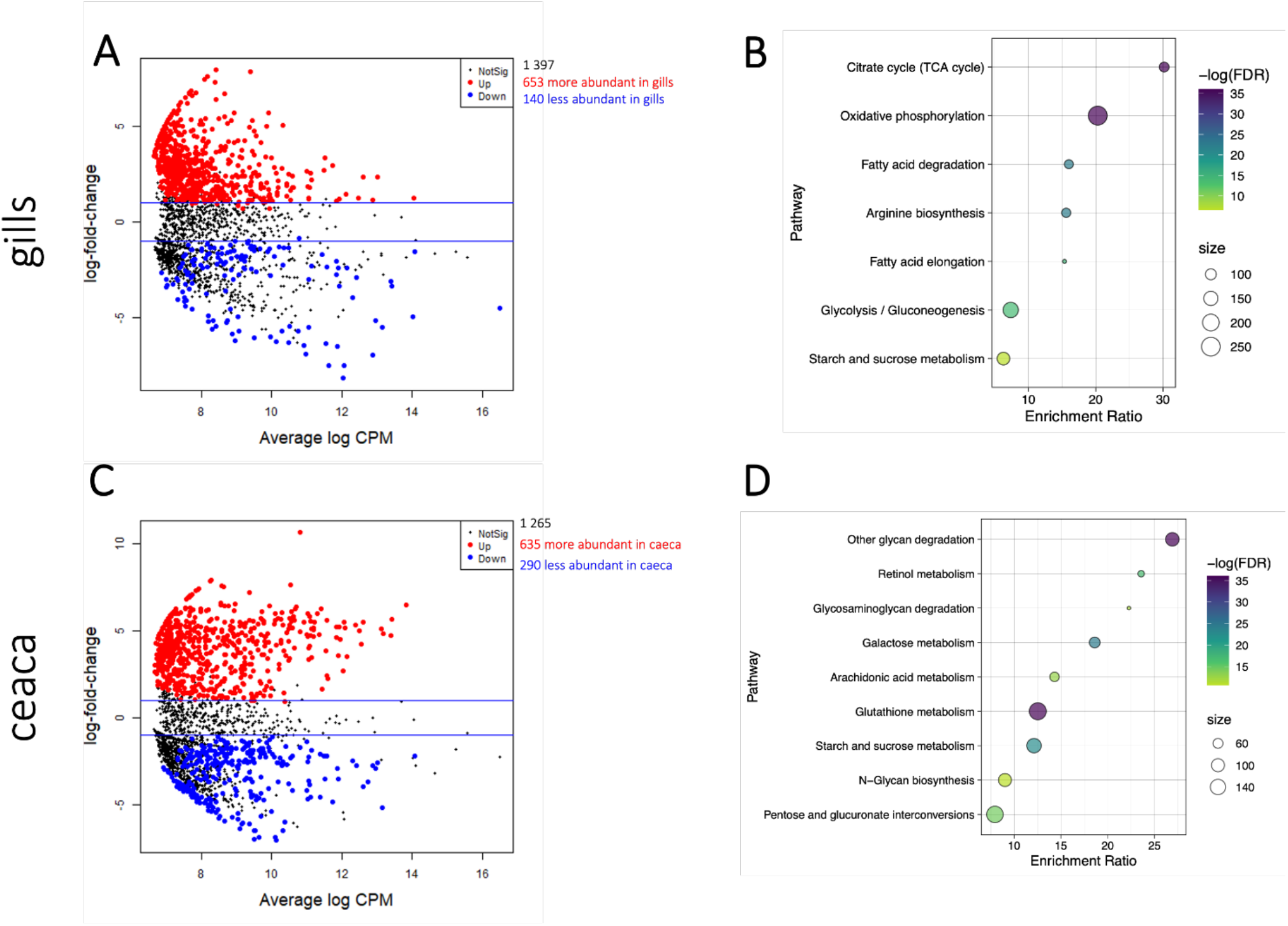
Differential protein abundances in gills and caeca of *G. fossarum* compared with the other organs (FDR < 0.05, LFC>2). (A) The MA plot, in red the most abundant proteins in gills, in blue the least abundant proteins in gills, (B) Enrichment plot for KEGG pathway categories in gills, the size of the dots is proportional to the number of genes present in the pathways, all the pathways presented have an FDR > 0.05, the more significant the enrichment, the darker the dot. (C) The MA plot in caeca. (D) Enrichment plot for KEGG pathway categories in the caeca.

In the caeca, 635 proteins were more abundant (FDR<0.05, FC>2) compared to other organs (Figure 6C, Table S2). Among these DE proteins, 228 were concomitantly annotated in GamfoCyc. Indeed, gammarid caeca have been previously identified as important organs for amino acids, lipids, carbohydrates, and vitamins (Leprêtre et al., 2023), but not specific pathways were described. The glycan degradation pathway was overrepresented in the caecum with an ER of 26.90% (FDR<2.2^e^-16) (Figure 6D, Table S6). This pathway has been reported to be involved in growth factor signaling and morphogenesis in arthropods (Scanlan et al., 2015) and the expression of genes involved in this pathway was altered by exposure to certain flame retardants (Scanlan et al., 2015). Proteins involved in the retinol metabolism pathway were also enriched in the caecum in the present study to around 23% (FDR=5.93e-08) (Table S6). Retinoids play crucial roles in many physiological processes, including embryonic development, morphogenesis, and cellular differentiation in vertebrates (Ghyselinck and Duester, 2019), and were also shown to be responsible for the development of crustaceans and insects (Liñán-Cabello et al., 2002; Nakamura et al., 2007). The retinoid system has been explored scarcely in crustaceans (Liñán-Cabello et al., 2002). However, some retinoids (oxidated forms of retinoate, retinoate isomers, and retinaldehyde isomers) were quantified in *G. fossarum* and were shown to fluctuate during the reproductive cycle in females and their levels were affected by methoprene, a juvenile hormone analog (Gauthier et al., 2023). Another over-represented KEGG pathway in the gammarid caeca was the glutathione metabolism with an ER of about 12% (FDR<2.2^e^-16) (Figure 6D, Table S6). Glutathione is known to be involved in detoxification, sequestering labile metals such as cadmium and preventing cytotoxicity (Khan et al., 2012). Recent studies by Gestin et al. (2023, 2022, 2021), have shown that gammarid caeca bioaccumulate and eliminate cadmium. These metabolic pathways may play a role in protecting against heavy metal exposure and oxidative stress following environmental variations.

Then we focused on the metabolic pathways present in the gonads. In total, 343 proteins were more abundant in the testes (FDR<0.05 and FC>2), while 187 DE proteins were overexpressed in the ovaries (FDR<0.05 and FC>2) (Figure S8A). Among these 530 DE proteins (male and female combined), 149 are annotated in GamfoCyc (Figure S8B).

In male gonads, glycolysis was the most enriched metabolic pathway with around 29% of enrichment ratio (ER) (FDR<2.2e-16) (Figure S8C and Table S3). Glycolysis is a pathway by which ATP molecules are formed from glucose to supply energy to the organism and has been highlighted as a key pathway in spermatogenesis in various vertebrate species. For instance, glycolysis has been shown to play an important role in early and late spermatogenesis in *Drosophila hydei* (Geer et al., 1972). Previous observations made using co-expression networks in gammarid male and female gonads showed that glycolysis was highly enriched in testes (Degli Esposti et al., 2019). Notably, through integrated metabolomics and transcriptomics analyses of *Macrobrachium nipponense* testes, it was found that glycolysis/gluconeogenesis and the tricarboxylic acid (TCA) cycle may play an essential role in promoting the process of male sexual differentiation and development by supplying ATP (Jin et al., 2020).

In the female gonads, metabolic pathways involved in glutathione and purine metabolism were found overrepresented with respectively around 11% and 10% of enrichment and an FDR of less than 0.05 (Figure S8D and Table S4). While predominant, amino acid metabolisms such as those involving alanine, aspartate, and glutamate were not significantly enriched. The glutathione system is involved in detoxification and oxidative stress response. Interestingly, in the crustacean *Metapenaeus ensis*, glutathione peroxidase was found to be specifically expressed in early ovaries, suggesting that the glutathione peroxidase might play a pivotal role in preventing oocytes from oxidative damage and thus in crustacean reproduction (Wu and Chu, 2010; Xia et al., 2013). Many aspects of oocyte maturation, such as mitosis and meiosis include DNA and RNA synthesis and rearrangements, and thus require an intense use purine nucleotides. Purines also act as enzyme cofactors, participate in cellular signaling, act as phosphate group donors to generate cellular energy (Carter et al., 2008).

## Conclusions

Our analysis demonstrates that transcriptomic data can be exploited by specific annotation systems such as CycADS to improve the annotation of metabolic pathways in species lacking genomic resources and the integration with proteomics data can contribute to improve the phenotypic characterization at the organ level and avoid potential biases (i.e., false negatives or false positives) coming from single omics approaches (Ge et al., 2003; Reeves et al., 2009). The reported results highlight the value of applying omics approaches on organs of small crustaceans to assess their metabolic specificities. Moreover, in the case of sentinel species used to assess the impact of contaminants on the ecosystems, this work put the basis to investigate the ability of environmental contaminants to disrupt metabolism depending on the target organ.

## Abbreviations

BLAST: Basic Local Alignment Search Tool
BUSCO: Benchmarking Universal Single-Copy Orthologues
CycADS: Cyc Annotation Database System
DE: Differentially expressed
EC: Enzyme Commission
FC: Fold Change
FDR: False Discovery Rate
GFF3: General Feature Format 3
*G. fossarum*: *Gammarus fossarum*
*GFBF / GFBM*: *Gammarus fossarum* female / male
GO: Gene Ontology
IsoPct: Isoform Percentage
KAAS: KEGG Automatic Annotation Server
KEGG: Kyoto Encyclopedia of Genes and Genomes
KO: KEGG Orthology
MDS: Multi-Dimensional Scaling
MS: Mass Spectrometry
NGS: Next Generation Sequencing
ORF: Open Reading Frame
PGDB: Pathway/Genome Database
ER: Enrichment Ratio
RNA-Seq: RNA Sequencing
RSEM: RNA-Seq by Expectation Maximisation
RT: Retention Time
SBML: Systems Biology Markup Language
SC: Spectral Count

## Data access

The sequences of the original reads are available in the “NCBI Sequence Reads Archive” under accession number SRR8089722 (“RNAseq GFBM,” 2018) (BioProject PRJNA497972 and BioSample SAMN10259937) for the male and under accession number SRR8089729 (“RNAseq GFBF,” 2018) (BioProject PRJNA497972 and BioSample SAMN10259936) for the female.

Original mass spectrometry data are available via the PRIDE repository with the dataset identifiers PXD040344. Peptide sequence data and spectral count data are available on https://doi.org/10.57745/HMQVCS.

All R scripts combining proteomic data integration, differential analysis, and metabolic pathway analysis are available on Github (https://github.com/NatachaKoenig/MultiomicsAnnotationGFB).

All functional annotations and metabolic network data are available on entrepot.recherche.data.gouv.fr (https://doi.org/10.57745/HMQVCS).

## Acknowledgements

The authors benefitted from the French GDR “Aquatic Ecotoxicology” framework which aims at fostering stimulating scientific discussions and collaborations for integrative approaches. This work has been supported by the APPROve project (ANR-18-CE34-0013-01) and by the Fédération de Recherche BioEEnVis (GamfoCyc project).

## Credit Author statement

NK: formal analysis, investigation, visualization, writing - original draft, writing - review & editing.

DDE: conceptualization, funding acquisition supervision, writing - original draft, writing - review & editing

OG: funding acquisition, review & editing

HQ, LG, ND, JCG, MK: methodology.

AL, PBP, ML and KS: formal analysis.

VN, AC, OG, ML, KS, HC, FC, SA, JA: writing - review & editing.

## Supplementary Information

**Table S1.**
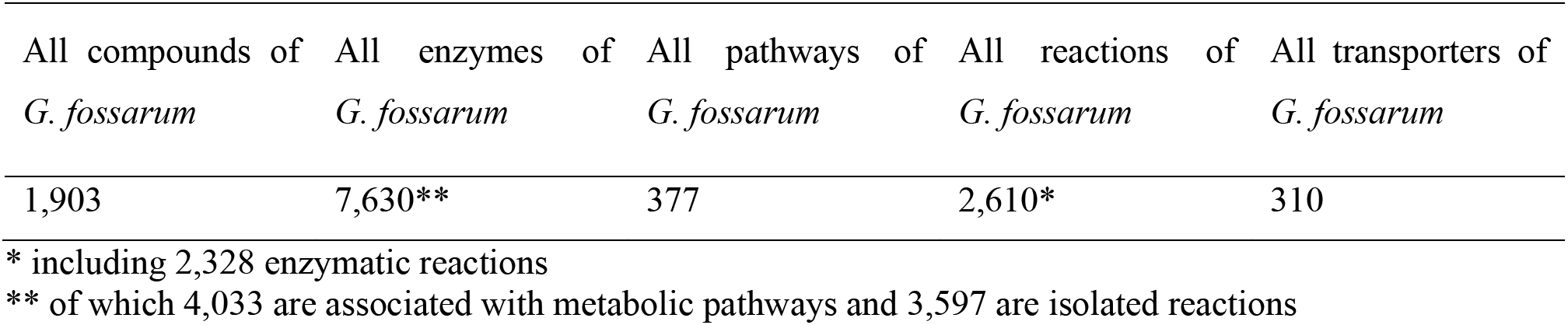
Summary table of GamfoCyc database statistics (extracted from the “Special SmartTables feature”)

Table S2. Differential expression analysis result tables for male gonads, female gonads, caeca, and gills in metabolic pathways in *Gammarus fossarum*.

Table S2 is provided as a separate Excel file with four tabs: differentially expressed (DE) proteins in male gonads, DE proteins in female gonads, DE proteins in gills, and DE proteins in caeca.

**Table S3.**
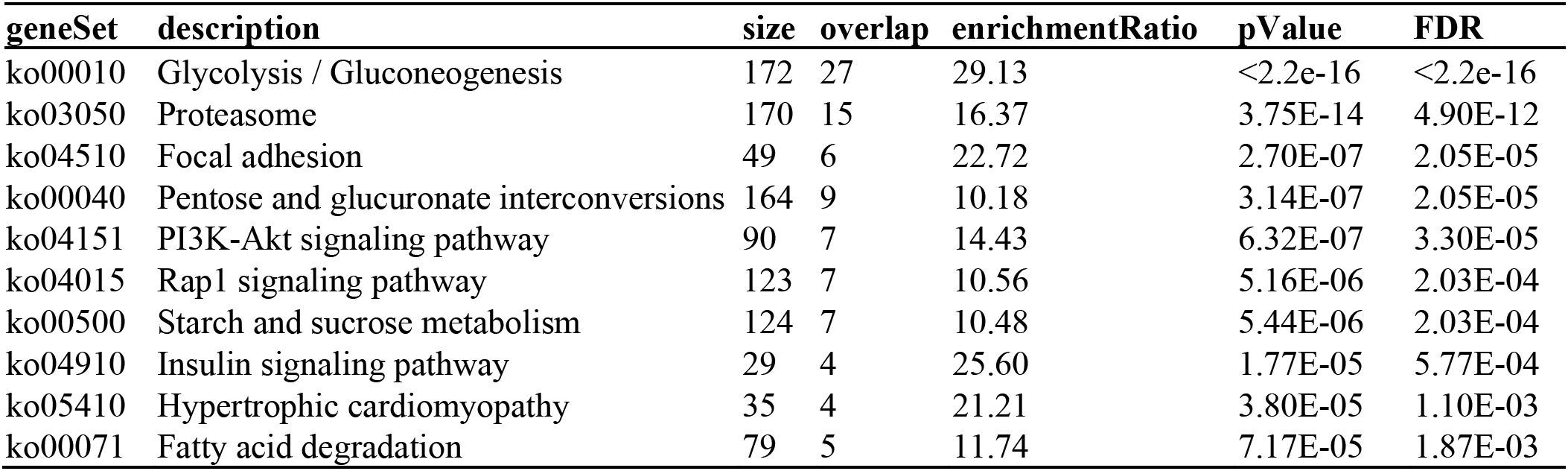
KEGG pathway enrichment analysis of the most abundant proteins in male gonads.

**Table S4.**
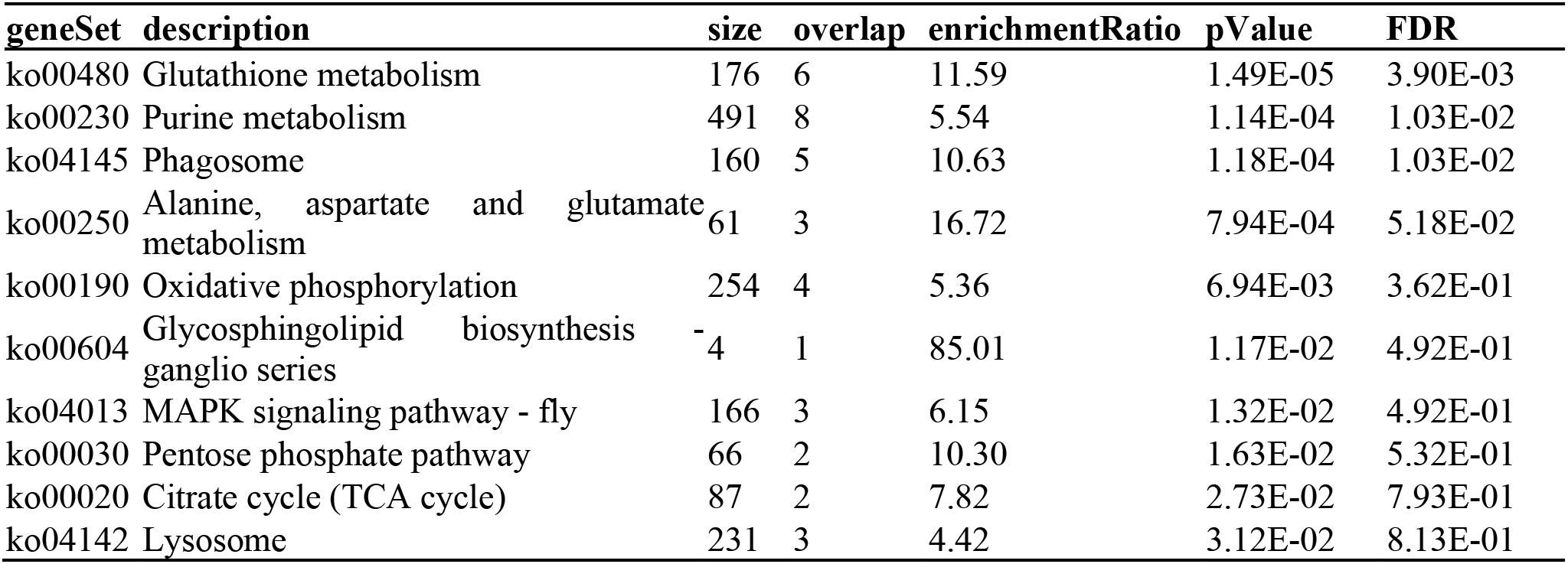
KEGG pathway enrichment analysis of the most abundant proteins in female gonads.

**Table S5.**
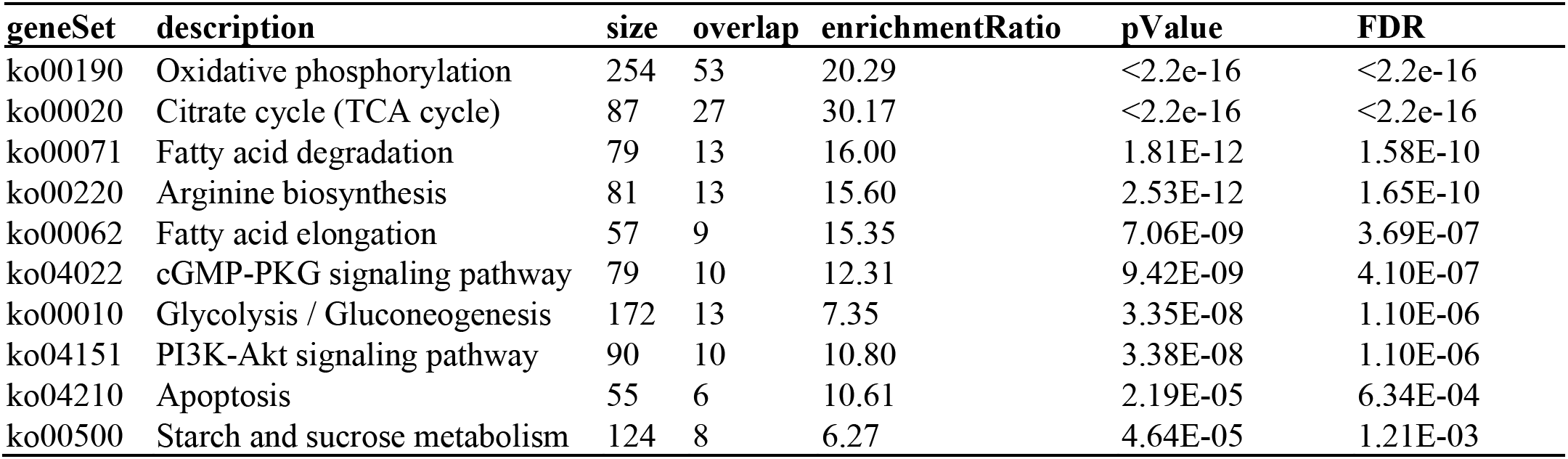
KEGG pathway enrichment analysis of the most abundant proteins in gills versus all other organs.

**Table S6.**
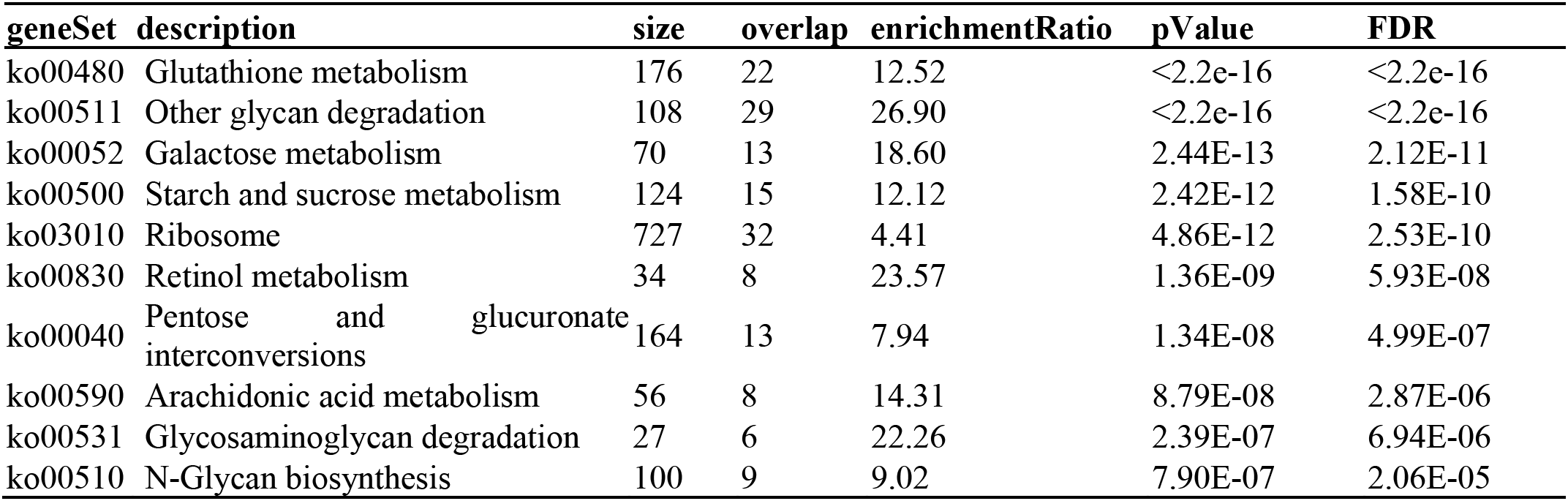
KEGG pathway enrichment analysis of the most abundant proteins in caeca versus all other organs.

**Figure S1.**
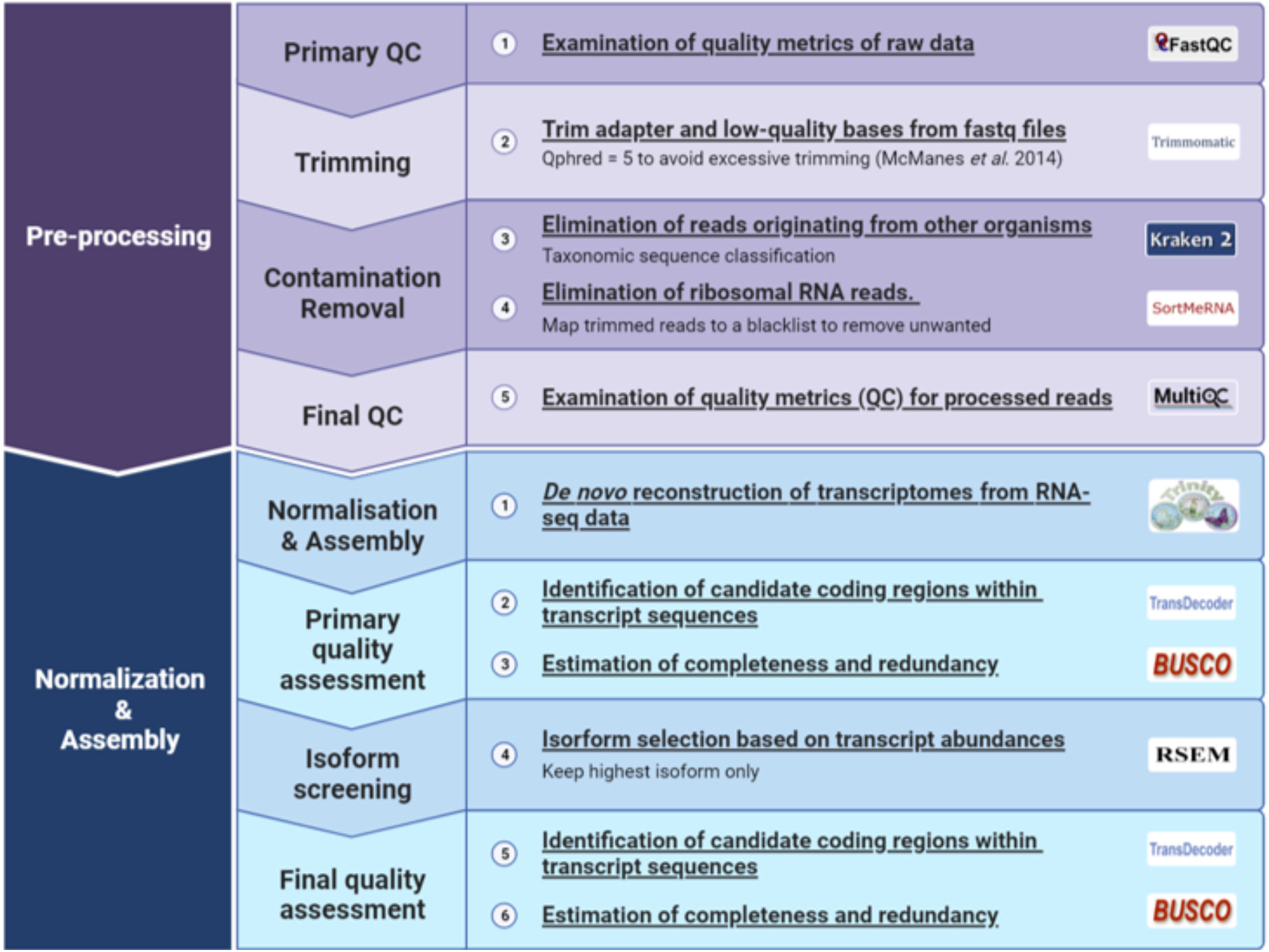
Reads pre-processing and de novo transcriptome assembly pipeline overview.

**Figure S2.**
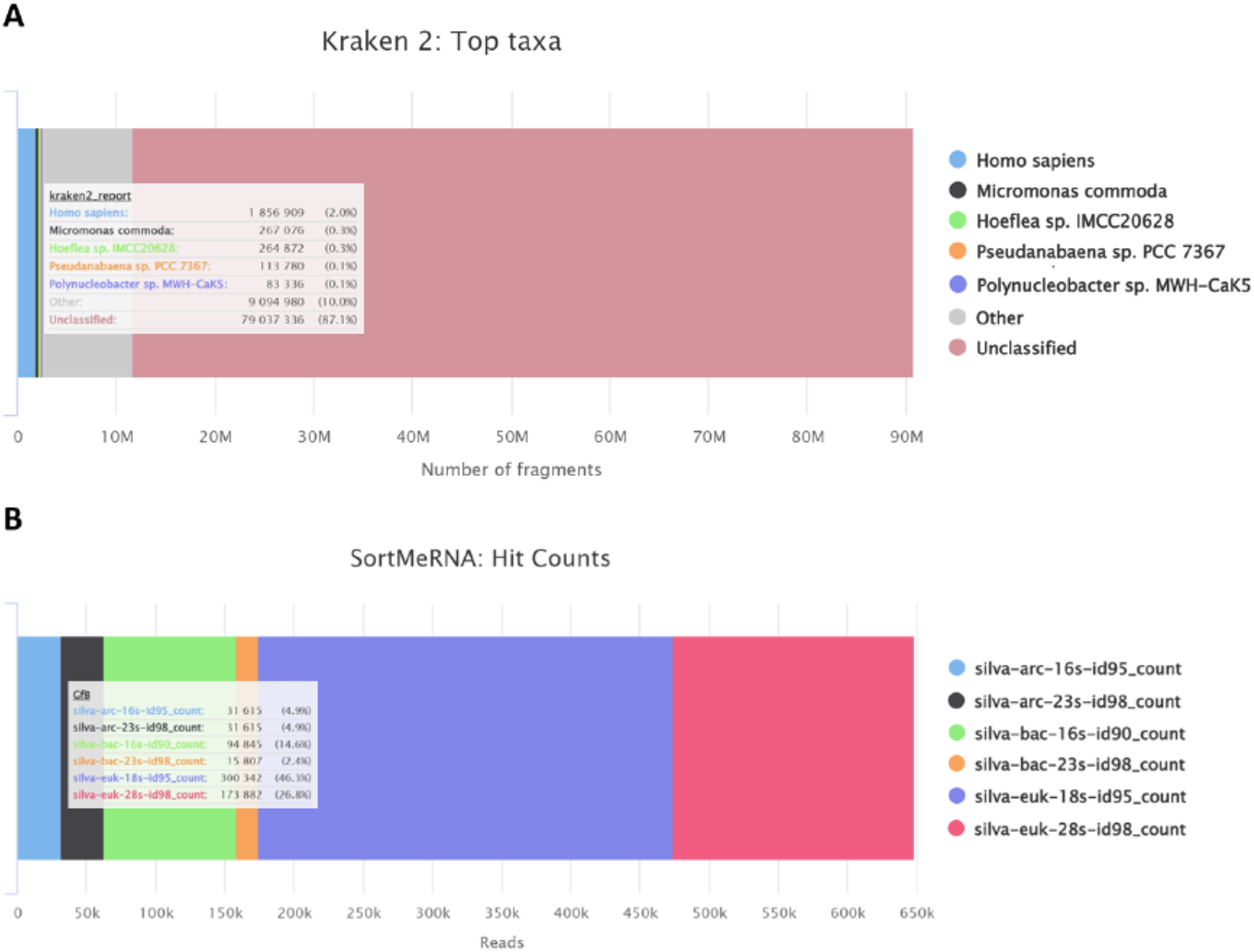
Barplots of the number of fragments/sequences classified for each taxon assigned by (A) Kraken2 and (B) SortMeRNA.

**Figure S3.**
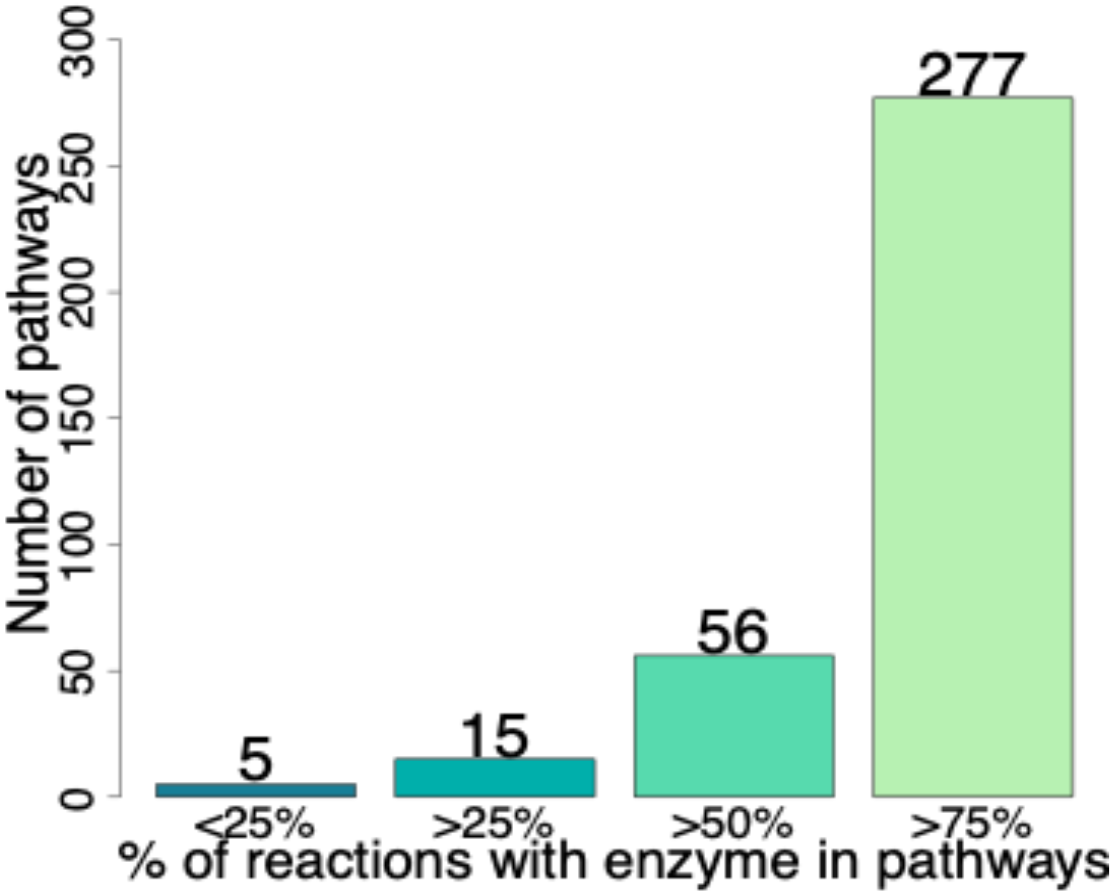
Completion of the global pathways of *Drosophila melanogaster* by MetExplore.

**Figure S4.**
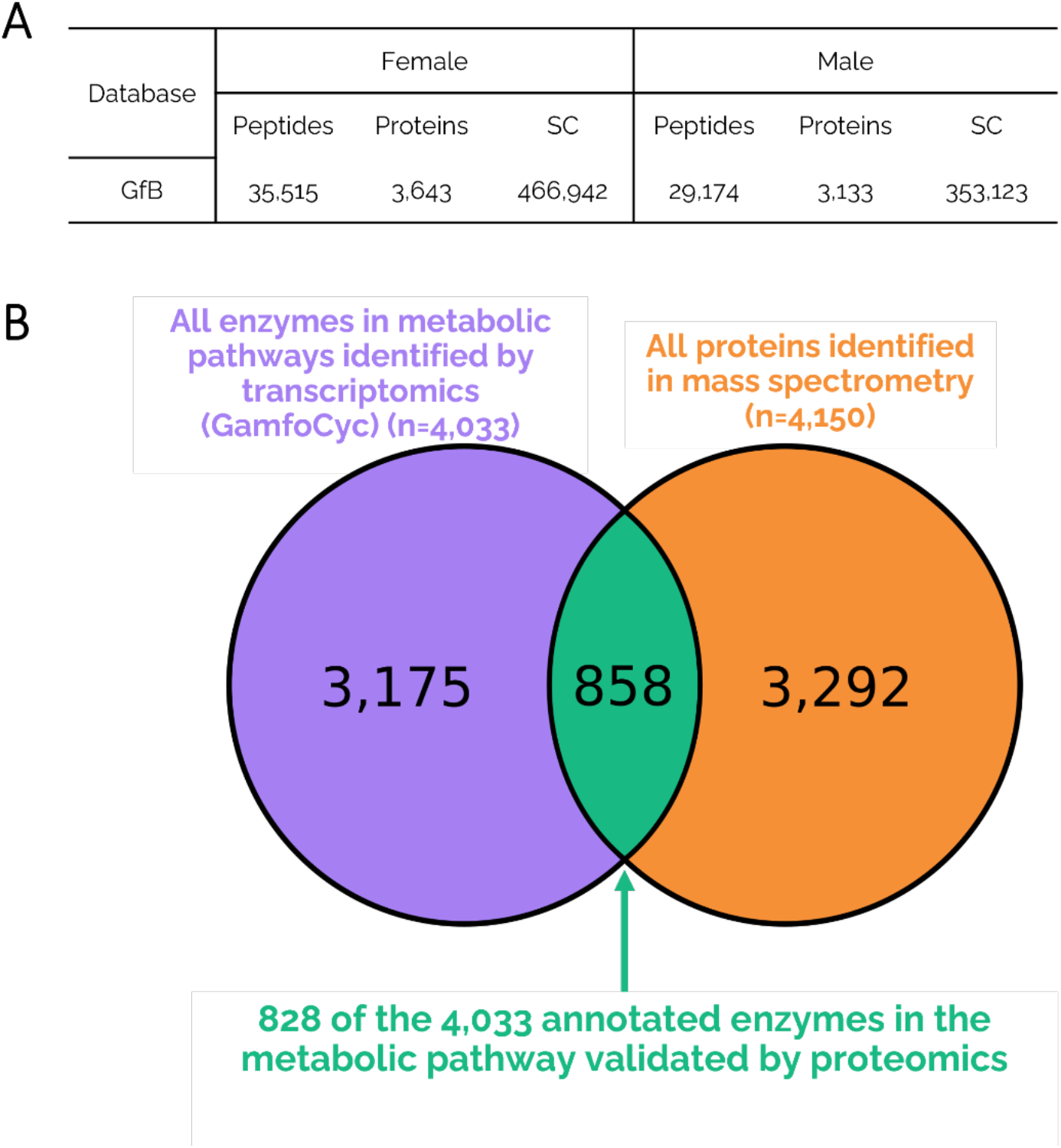
Integration of proteomic data for metabolic annotation (A) Mass spectrometry results interpreted with GfB transcriptome-derived database, SC: spectral count (B) Venn diagram of Gammarus fossarum enzymes annotated by CycADS and validated using proteomic.

**Figure S5.**
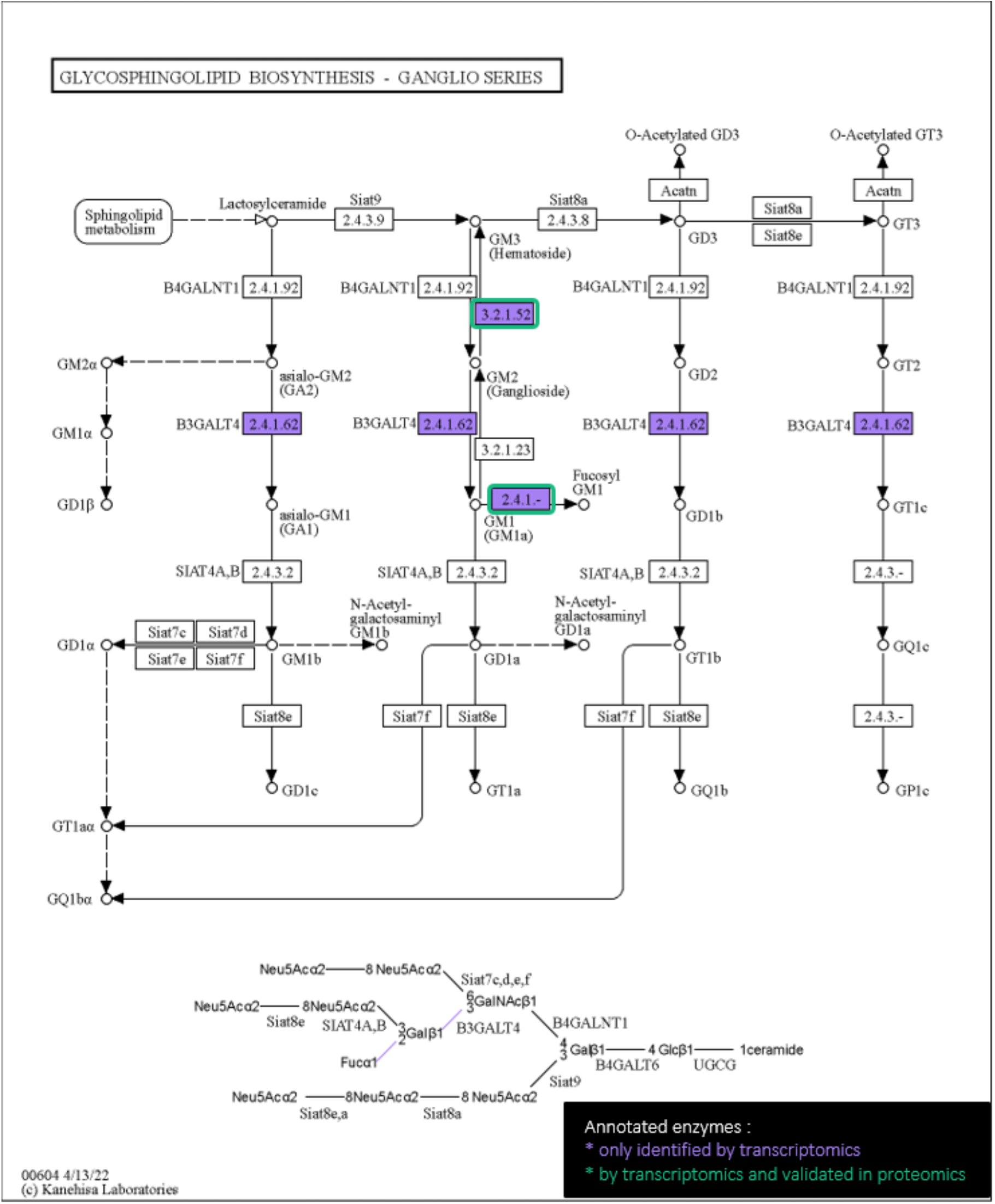
Example of mapped enzymes of *Gammarus fossarum* involved in the biosynthesis of glycosphingolipid pathway. Pathway module (functional unit of gene sets in metabolic pathway) is highlighted in green when the correspondent EC was annotated by transcriptomics and validated in proteomics and highlight in purple when the correspondent EC was annotated only by transcriptomics.

**Figure S6.**
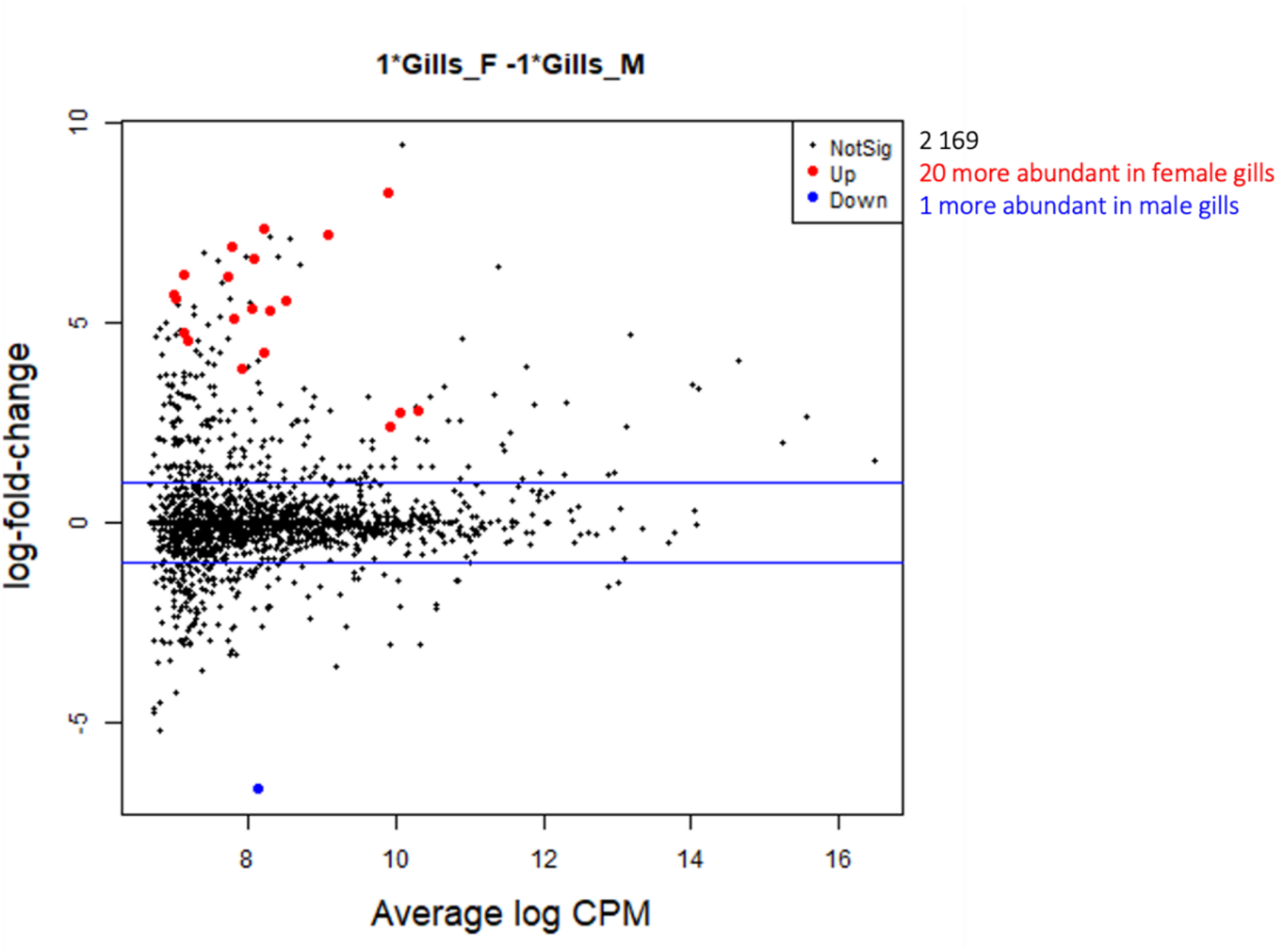
MA plot of differential analysis of female versus male gills

**Figure S7.**
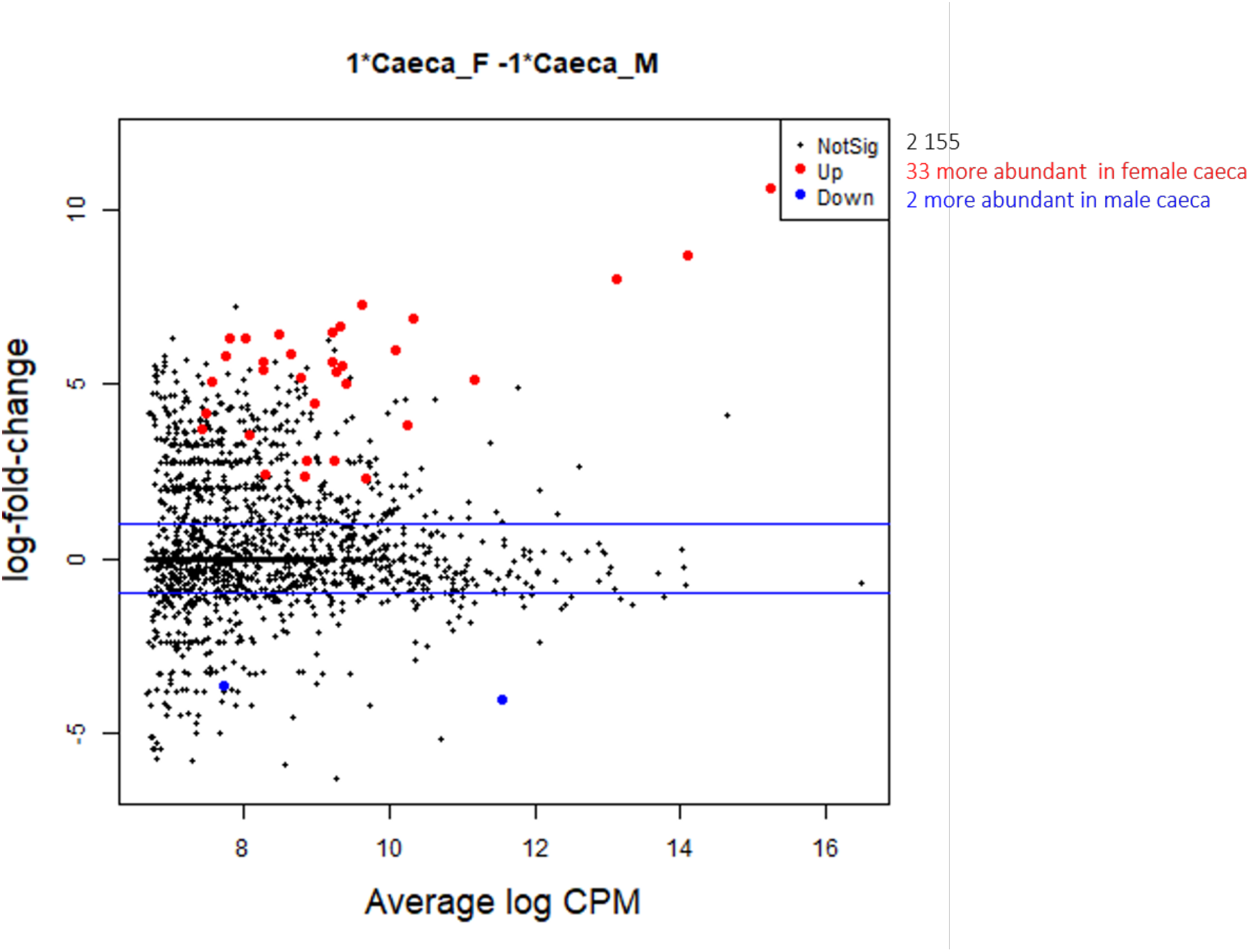
MA plot of differential analysis of male versus female caeca

**Figure S8.**
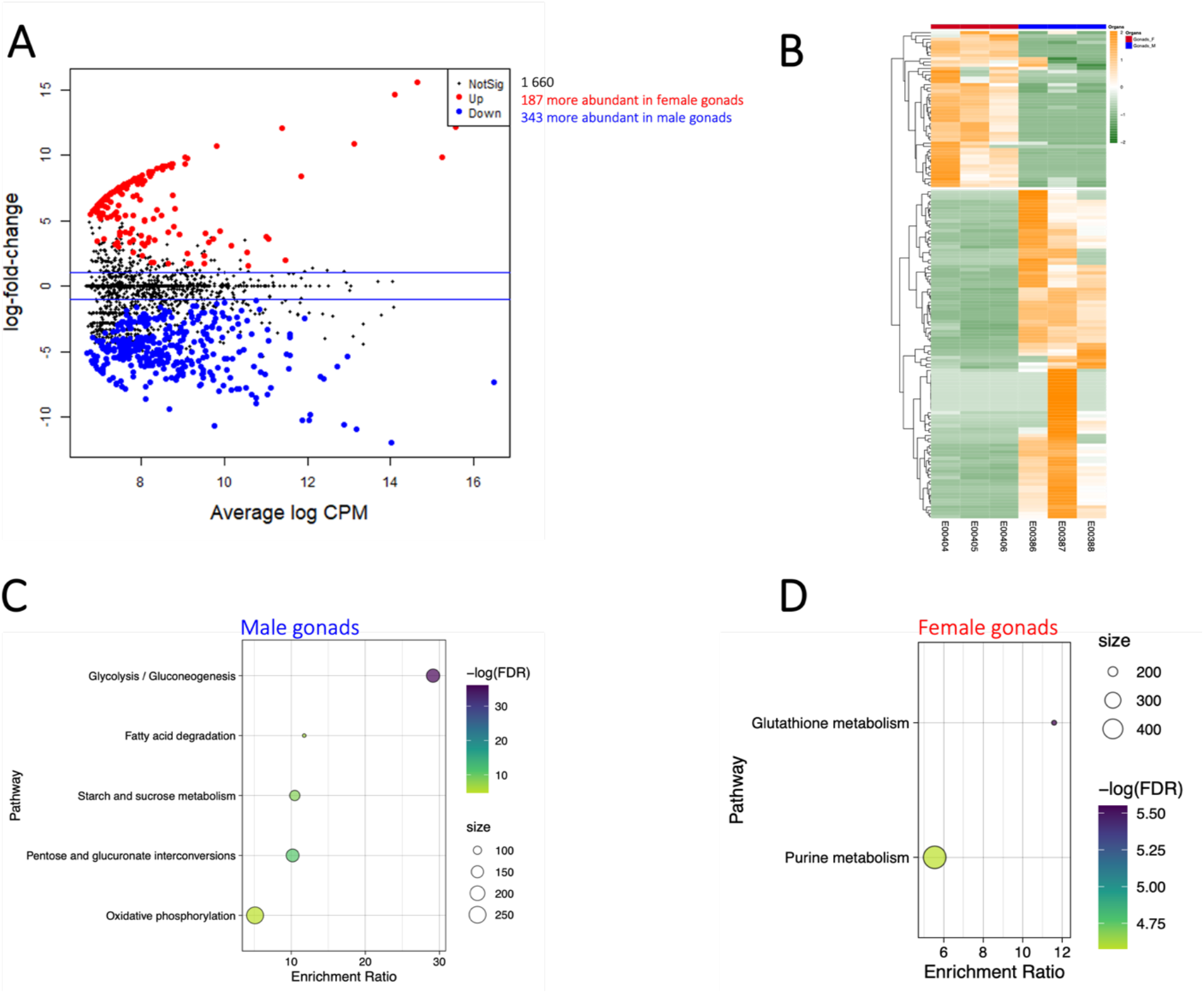
Differential protein abundances between gammarid gonads (FDR < 0.05, LFC > 2). (A) The MA plot, in red the most abundant proteins in female gonads, in blue the most abundant proteins in male gonads, (B) the heatmap of significantly abundant proteins validated by proteomics in the female gonads (red) and male gonads (blue). KEGG pathways enrichment plot for male gonads (C) and female gonads (D), the size of the dots is proportional to the number of genes present in the pathways, all the pathways presented have an FDR > 0.05, the more significant the enrichment, the darker the dot.

