## Supplementary Information for "Proteogenomic reconstruction of organ-specific metabolic networks in an environmental sentinel species, the amphipod *Gammarus fossarum*"

1 INRAE, UR RiverLy, Ecotoxicology Team. Centre de Lyon-Grenoble Auvergne Rhône Alpes, 5 rue de la Doua CS 20244, 69625 Villeurbanne, France

2 INRAE, INSA Lyon, BF2I, UMR203, 69621 Villeurbanne, France

3 PRABI, Rhône-Alpes Bioinformatics Center, Université Lyon 1, Villeurbanne, France, UMS 3601, Institut Français de Bioinformatique, IFB-Core, Évry, France.

4 Université Paris-Saclay, Département Médicaments et Technologies pour la Santé (DMTS), CEA, INRAE, SPI-Li2D, F-30207 Bagnols-sur-Céze, France

5 University of Lyon, CNRS, Institut des Sciences Analytiques, UMR 5280, 5 rue de la Doua, F-69100 Villeurbanne, France

[#]Corresponding Author: Davide Degli Esposti, INRAE, UR RiverLy, Ecotoxicology Team. Centre de Lyon-Grenoble Auvergne Rhône Alpes, 5 rue de la Doua CS 20244, 69625 Villeurbanne, France.

### Supplemental material.

9 pages, 6 tables, 9 figures.

#### Table of Contents

##### Tables

**Table S1. Summary table of GamfoCyc database statistics (extracted from the "Special SmartTables feature")** 4

**Table S2: The differential expression analysis result tables for male gonads, female gonads, caeca, and gills in metabolic pathways in *Gammarus fossarum*.** 4

**Table S3. KEGG pathway enrichment analysis of the most abundant proteins in male gonads.** 4

**Table S4. KEGG pathway enrichment analysis of the most abundant proteins in female gonads.** 4

**Table S5. KEGG pathway enrichment analysis of the most abundant proteins in gills versus all other organs.** 5

**Table S6. KEGG pathway enrichment analysis of the most abundant proteins in caeca versus all other organs.** 5

##### Figures

**Figure S1. Reads pre-processing and de novo transcriptome assembly pipeline overview.** 6

**Figure S2. Barplots of the number of fragments/sequences classified for each taxon assigned by (A) Kraken2 and (B) SortMeRNA.** 7

**Figure S3. Completion of the global pathways of *Drosophila melanogaster* by MetExplore.** 7

**Figure S4. Integration of proteomic data for metabolic annotation (A) Mass spectrometry results interpreted with GfB transcriptome-derived database, SC: spectral count (B) Venn diagram of Gammarus fossarum enzymes annotated by CycADS and validated using proteomic.** 8

**Figure S5. Example of mapped enzymes of *Gammarus fossarum* involved in the biosynthesis of glycosphingolipid pathway. Pathway module (functional unit of gene sets in metabolic pathway) is highlighted in green when the correspondent EC was annotated by transcriptomics and validated in proteomics and highlight in purple when the correspondent EC was annotated only by transcriptomics.** 9

**Figure S6. MA plot of differential analysis of female versus male gills** 10

**Figure S7. MA plot of differential analysis of male versus female caeca** 11

**Figure S8. Differential protein abundances between gammarid gonads (FDR < 0.05, LFC > 2). (A) The MA plot, in red the most abundant proteins in female gonads, in blue the most abundant proteins in male gonads, (B) the heatmap of significantly abundant proteins validated by proteomics in the female gonads (red) and male gonads (blue). KEGG pathways enrichment plot for male gonads (C) and female gonads (D), the size of the dots is proportional to the number of genes present in the pathways, all the pathways presented have an FDR > 0.05, the more significant the enrichment, the darker the dot. 12**

Table S1. Summary table of GamfoCyc database statistics (extracted from the “Special SmartTables feature”)

| All compounds of *G. fossarum* | All enzymes of *G. fossarum* | All pathways of *G. fossarum* | All reactions of *G. fossarum* | All transporters of *G. fossarum* |
|---|---|---|---|---|
| 1,903 | 7,630** | 377 | 2,610* | 310 |

* including 2,328 enzymatic reactions
** of which 4,033 are associated with metabolic pathways and 3,597 are isolated reactions

Table S2: The differential expression analysis result tables for male gonads, female gonads, caeca, and gills in metabolic pathways in *Gammarus fossarum*.

Table S2 is provided as a separate Excel file with four tabs: differentially expressed (DE) proteins in male gonads, DE proteins in female gonads, DE proteins in gills, and DE proteins in caeca.

Table S3. KEGG pathway enrichment analysis of the most abundant proteins in male gonads.

| geneSet | description | size | overlap | enrichmentRatio | pValue | FDR |
|---|---|---|---|---|---|---|
| ko00010 | Glycolysis / Gluconeogenesis | 172 | 27 | 29.13 | <2.2e-16 | <2.2e-16 |
| ko03050 | Proteasome | 170 | 15 | 16.37 | 3.75E-14 | 4.90E-12 |
| ko04510 | Focal adhesion | 49 | 6 | 22.72 | 2.70E-07 | 2.05E-05 |
| ko00040 | Pentose and glucuronate interconversions | 164 | 9 | 10.18 | 3.14E-07 | 2.05E-05 |
| ko04151 | PI3K-Akt signaling pathway | 90 | 7 | 14.43 | 6.32E-07 | 3.30E-05 |
| ko04015 | Rap1 signaling pathway | 123 | 7 | 10.56 | 5.16E-06 | 2.03E-04 |
| ko00500 | Starch and sucrose metabolism | 124 | 7 | 10.48 | 5.44E-06 | 2.03E-04 |
| ko04910 | Insulin signaling pathway | 29 | 4 | 25.60 | 1.77E-05 | 5.77E-04 |
| ko05410 | Hypertrophic cardiomyopathy | 35 | 4 | 21.21 | 3.80E-05 | 1.10E-03 |
| ko00071 | Fatty acid degradation | 79 | 5 | 11.74 | 7.17E-05 | 1.87E-03 |

Table S4. KEGG pathway enrichment analysis of the most abundant proteins in female gonads.

| geneSet | description | size | overlap | enrichmentRatio | pValue | FDR |
|---|---|---|---|---|---|---|
| ko00480 | Glutathione metabolism | 176 | 6 | 11.59 | 1.49E-05 | 3.90E-03 |
| ko00230 | Purine metabolism | 491 | 8 | 5.54 | 1.14E-04 | 1.03E-02 |
| ko04145 | Phagosome | 160 | 5 | 10.63 | 1.18E-04 | 1.03E-02 |
| ko00250 | Alanine, aspartate and glutamate metabolism | 61 | 3 | 16.72 | 7.94E-04 | 5.18E-02 |
| ko00190 | Oxidative phosphorylation | 254 | 4 | 5.36 | 6.94E-03 | 3.62E-01 |
| ko00604 | Glycosphingolipid biosynthesis - ganglio series | 4 | 1 | 85.01 | 1.17E-02 | 4.92E-01 |
| ko04013 | MAPK signaling pathway - fly | 166 | 3 | 6.15 | 1.32E-02 | 4.92E-01 |
| ko00030 | Pentose phosphate pathway | 66 | 2 | 10.30 | 1.63E-02 | 5.32E-01 |
| ko00020 | Citrate cycle (TCA cycle) | 87 | 2 | 7.82 | 2.73E-02 | 7.93E-01 |
| ko04142 | Lysosome | 231 | 3 | 4.42 | 3.12E-02 | 8.13E-01 |

Table S5. KEGG pathway enrichment analysis of the most abundant proteins in gills versus all other organs.

| geneSet | description | size | overlap | enrichmentRatio | pValue | FDR |
|---|---|---|---|---|---|---|
| ko00190 | Oxidative phosphorylation | 254 | 53 | 20.29 | <2.2e-16 | <2.2e-16 |
| ko00020 | Citrate cycle (TCA cycle) | 87 | 27 | 30.17 | <2.2e-16 | <2.2e-16 |
| ko00071 | Fatty acid degradation | 79 | 13 | 16.00 | 1.81E-12 | 1.58E-10 |
| ko00220 | Arginine biosynthesis | 81 | 13 | 15.60 | 2.53E-12 | 1.65E-10 |
| ko00062 | Fatty acid elongation | 57 | 9 | 15.35 | 7.06E-09 | 3.69E-07 |
| ko04022 | cGMP-PKG signaling pathway | 79 | 10 | 12.31 | 9.42E-09 | 4.10E-07 |
| ko00010 | Glycolysis / Gluconeogenesis | 172 | 13 | 7.35 | 3.35E-08 | 1.10E-06 |
| ko04151 | PI3K-Akt signaling pathway | 90 | 10 | 10.80 | 3.38E-08 | 1.10E-06 |
| ko04210 | Apoptosis | 55 | 6 | 10.61 | 2.19E-05 | 6.34E-04 |
| ko00500 | Starch and sucrose metabolism | 124 | 8 | 6.27 | 4.64E-05 | 1.21E-03 |

Table S6. KEGG pathway enrichment analysis of the most abundant proteins in caeca versus all other organs.

| geneSet | description | size | overlap | enrichmentRatio | pValue | FDR |
|---|---|---|---|---|---|---|
| ko00480 | Glutathione metabolism | 176 | 22 | 12.52 | <2.2e-16 | <2.2e-16 |
| ko00511 | Other glycan degradation | 108 | 29 | 26.90 | <2.2e-16 | <2.2e-16 |
| ko00052 | Galactose metabolism | 70 | 13 | 18.60 | 2.44E-13 | 2.12E-11 |
| ko00500 | Starch and sucrose metabolism | 124 | 15 | 12.12 | 2.42E-12 | 1.58E-10 |
| ko03010 | Ribosome | 727 | 32 | 4.41 | 4.86E-12 | 2.53E-10 |
| ko00830 | Retinol metabolism | 34 | 8 | 23.57 | 1.36E-09 | 5.93E-08 |
| ko00040 | Pentose and glucuronate interconversions | 164 | 13 | 7.94 | 1.34E-08 | 4.99E-07 |
| ko00590 | Arachidonic acid metabolism | 56 | 8 | 14.31 | 8.79E-08 | 2.87E-06 |
| ko00531 | Glycosaminoglycan degradation | 27 | 6 | 22.26 | 2.39E-07 | 6.94E-06 |
| ko00510 | N-Glycan biosynthesis | 100 | 9 | 9.02 | 7.90E-07 | 2.06E-05 |

Figure S1. Reads pre-processing and de novo transcriptome assembly pipeline overview.

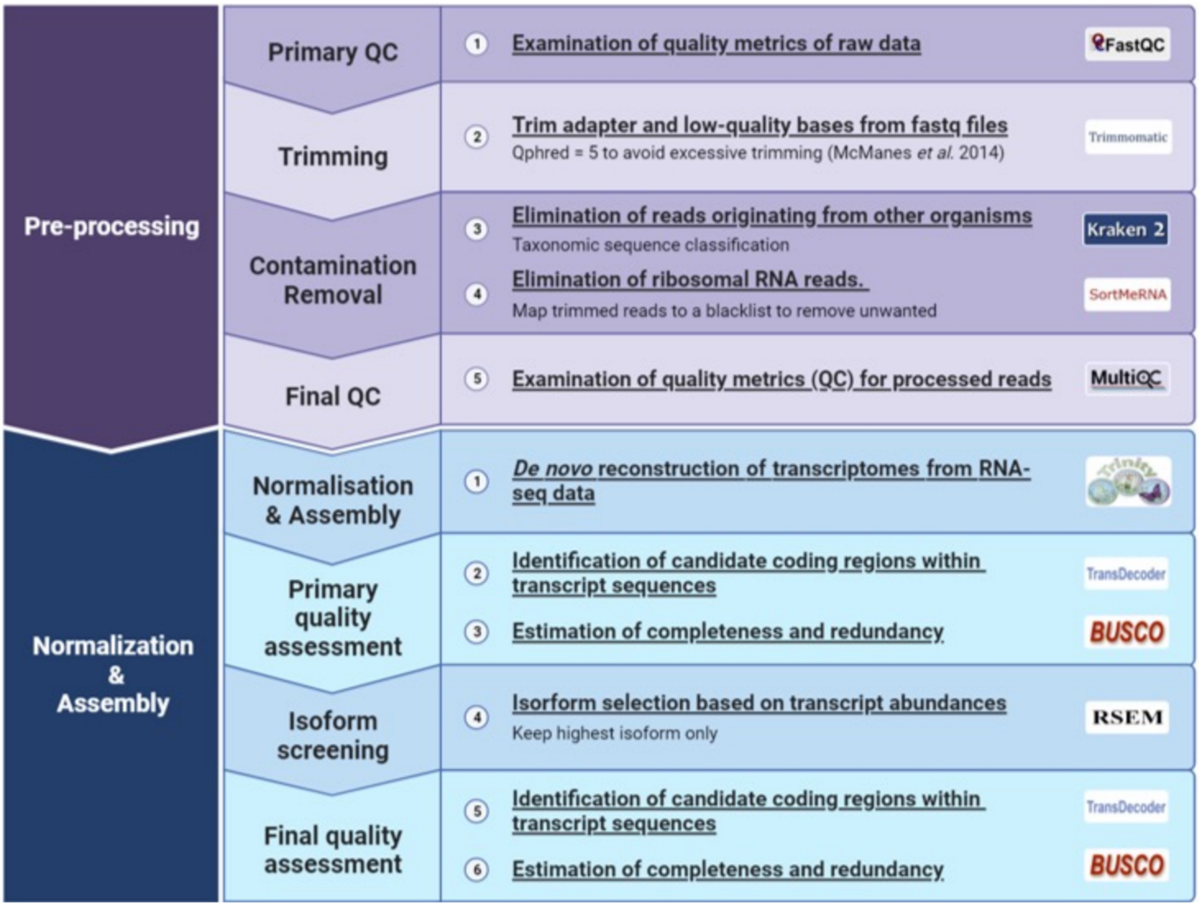

Figure S2. Barplots of the number of fragments/sequences classified for each taxon assigned by (A) Kraken2 and (B) SortMeRNA.

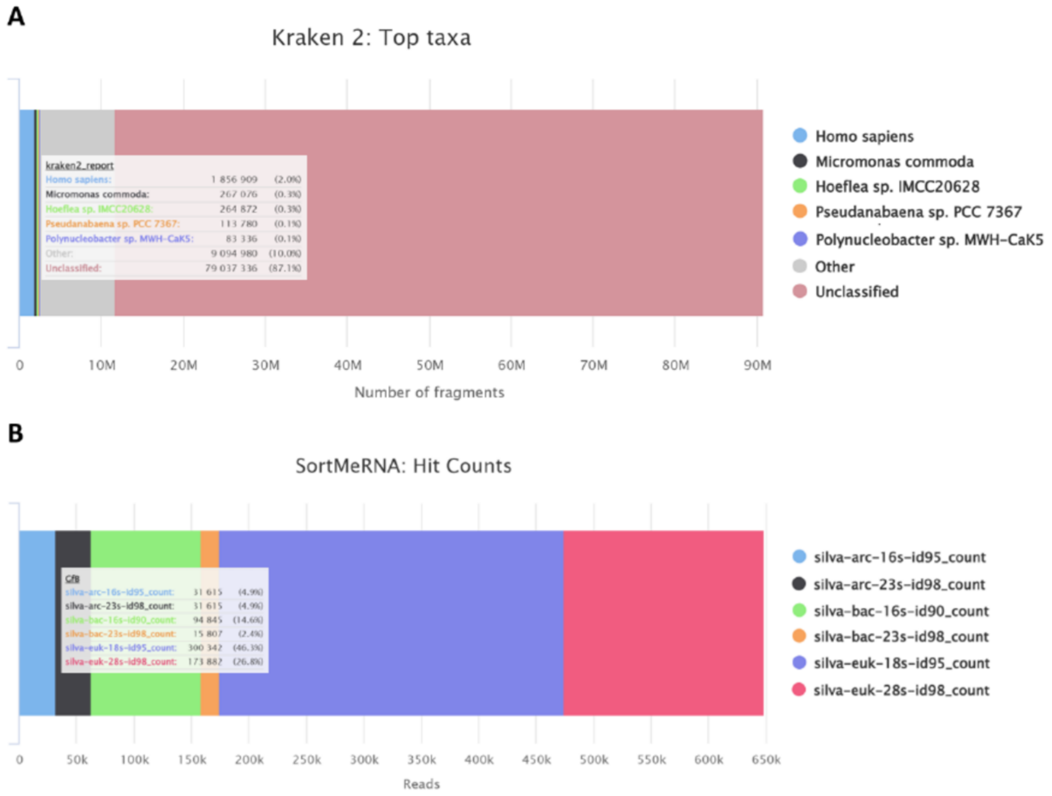


Figure S3. Completion of the global pathways of *Drosophila melanogaster* by MetExplore.

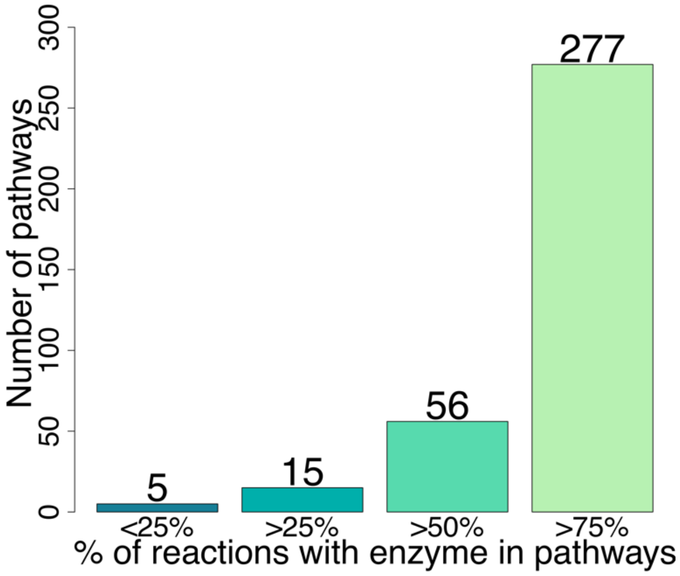

Figure S4. Integration of proteomic data for metabolic annotation (A) Mass spectrometry results interpreted with GfB transcriptome-derived database, SC: spectral count (B) Venn diagram of Gammarus fossarum enzymes annotated by CycADS and validated using proteomic.

A

| Database | Female | | | Male | | |
|---|---|---|---|---|---|---|
| | Peptides | Proteins | SC | Peptides | Proteins | SC |
| GfB | 35.515 | 3.643 | 466.942 | 29.174 | 3.133 | 353.123 |

B

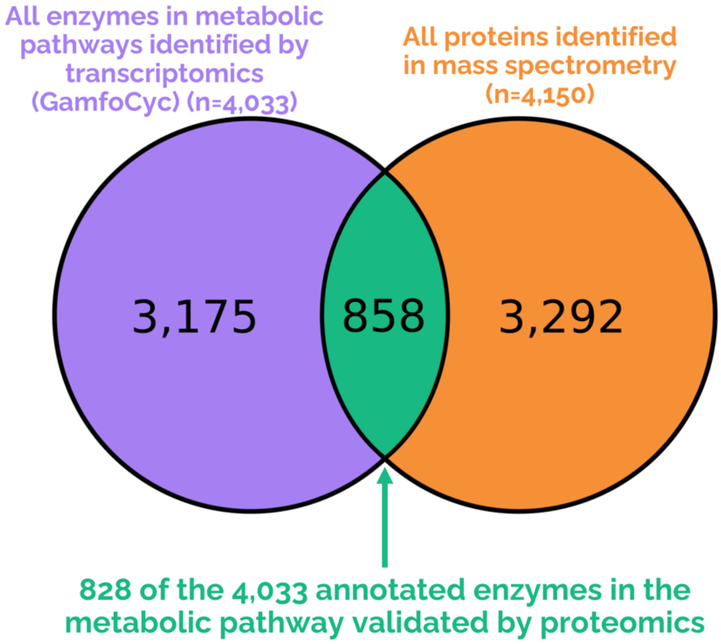

Figure S5. Example of mapped enzymes of *Gammarus fossarum* involved in the biosynthesis of glycosphingolipid pathway. Pathway module (functional unit of gene sets in metabolic pathway) is highlighted in green when the correspondent EC was annotated by transcriptomics and validated in proteomics and highlight in purple when the correspondent EC was annotated only by transcriptomics.

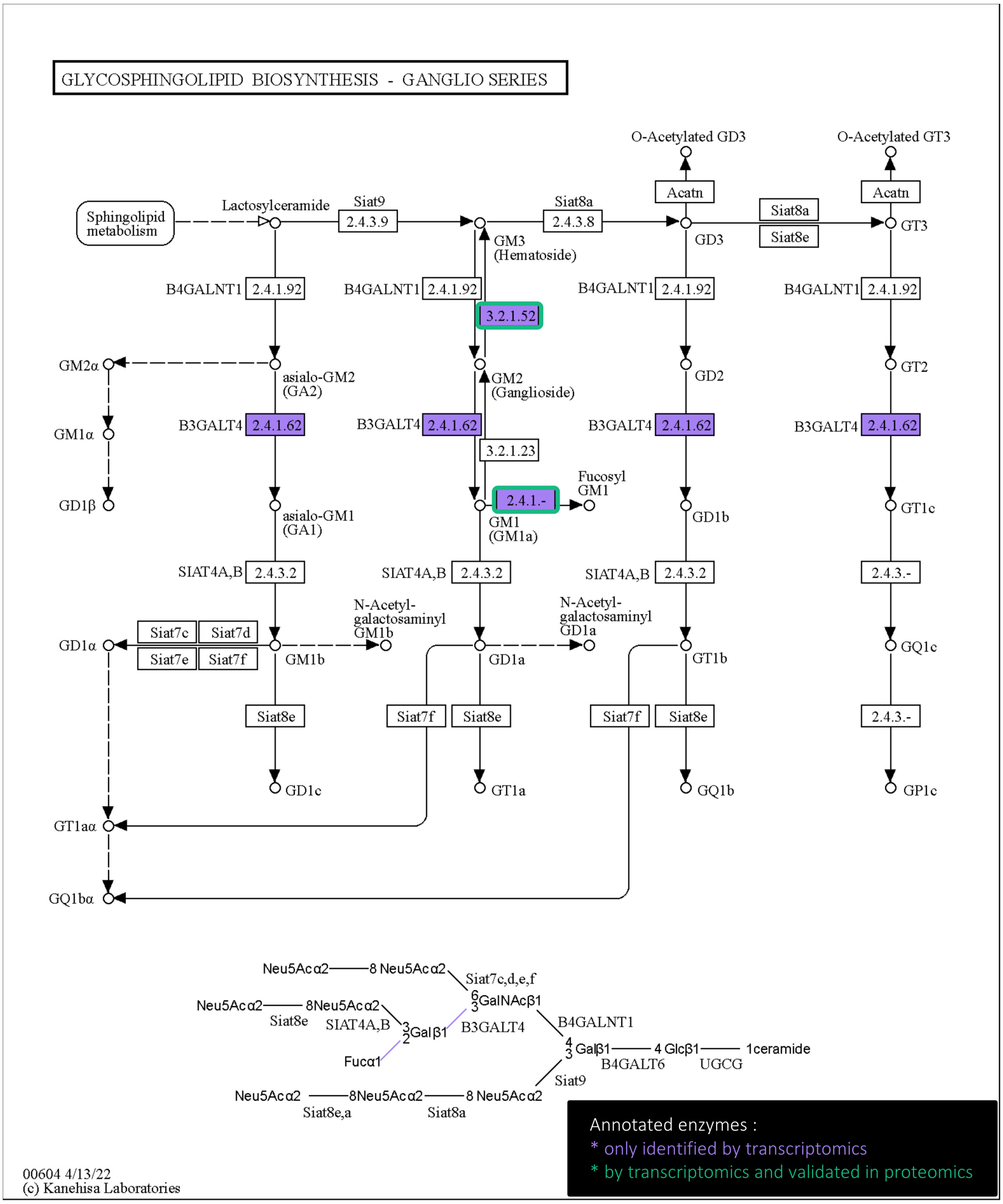

Figure S6. MA plot of differential analysis of female versus male gills

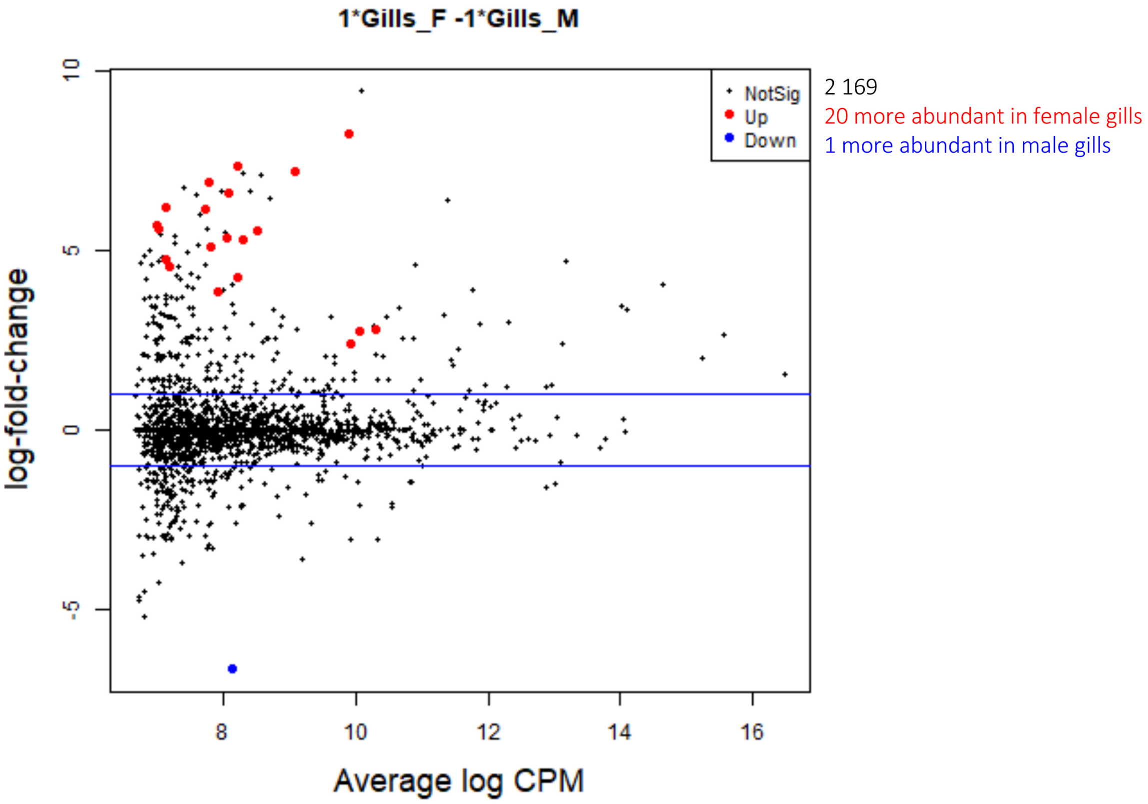

Figure S7. MA plot of differential analysis of male versus female caeca

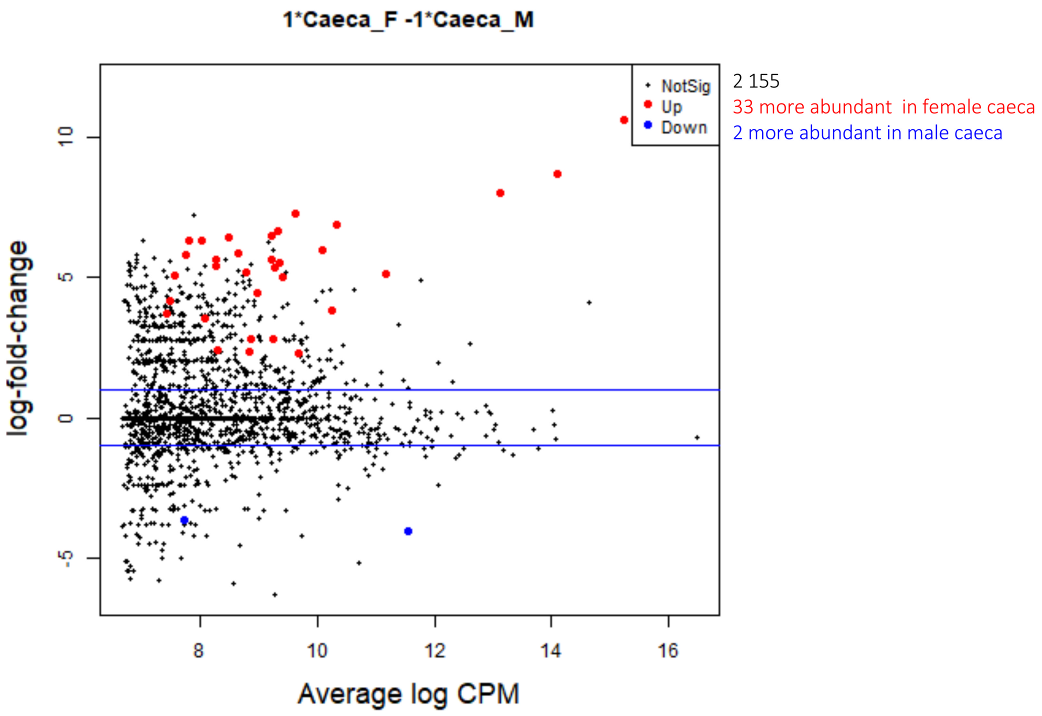

Figure S8. Differential protein abundances between gammarid gonads (FDR < 0.05, LFC > 2). (A) The MA plot, in red the most abundant proteins in female gonads, in blue the most abundant proteins in male gonads, (B) the heatmap of significantly abundant proteins validated by proteomics in the female gonads (red) and male gonads (blue). KEGG pathways enrichment plot for male gonads (C) and female gonads (D), the size of the dots is proportional to the number of genes present in the pathways, all the pathways presented have an FDR > 0.05, the more significant the enrichment, the darker the dot.

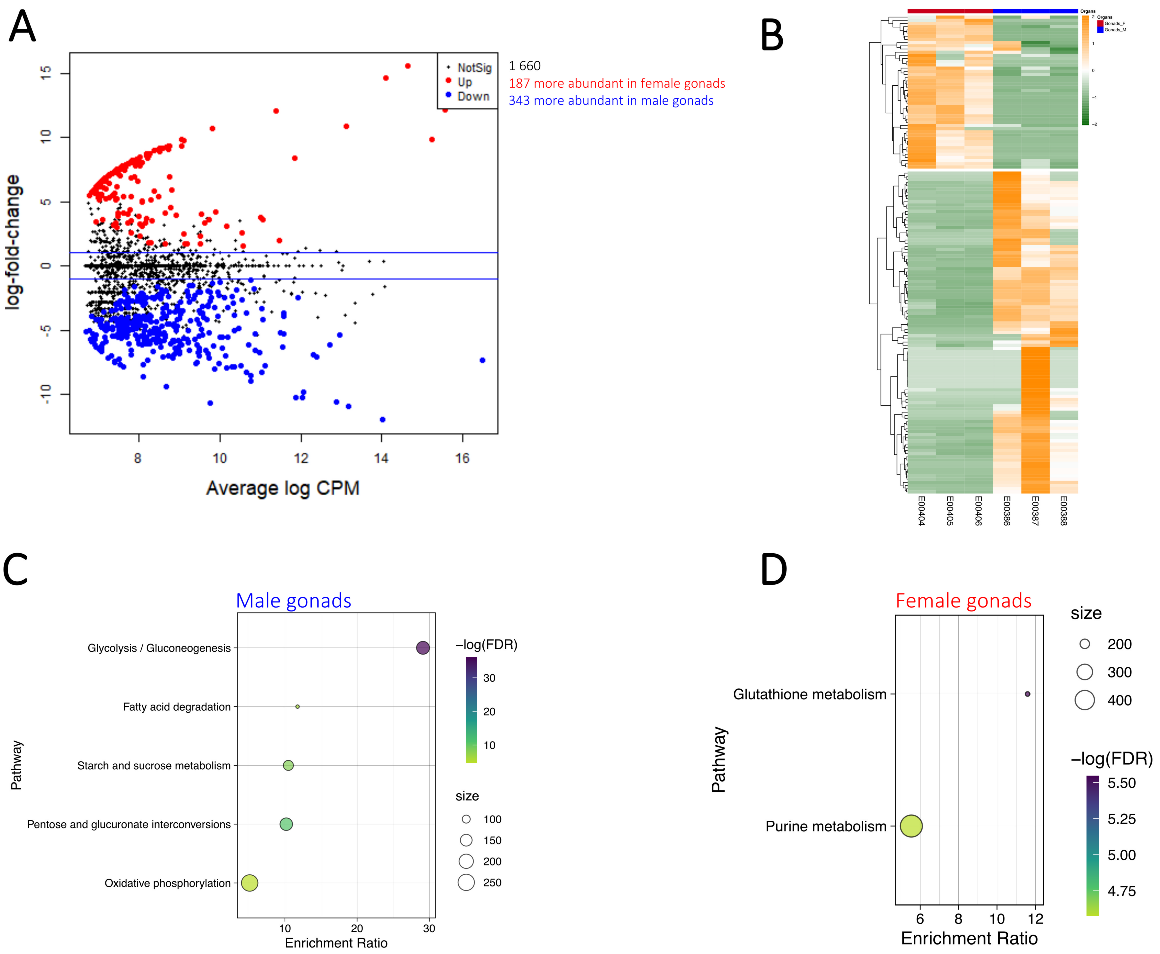
